# Axial Elongation of Caudalized Human Organoids Mimics Neural Tube Development

**DOI:** 10.1101/2020.03.05.979732

**Authors:** ARG Libby, DA Joy, NH Elder, EA Bulger, MZ Krakora, EA Gaylord, F Mendoza-Camacho, JC Butts, TC McDevitt

## Abstract

Axial elongation of the neural tube is critical during mammalian embryogenesis to establish the anterior-posterior body axis^1^, but this process is difficult to interrogate directly because it occurs post-implantation^2,3^. Here we report an organoid model of neural tube extension by caudalized human pluripotent stem cell (hPSC) aggregates that recapitulates the morphologic and temporal gene expression patterns of neural tube development. Axially extending organoids consisting largely of longitudinally elongated neuroepithelial compartments also contained TBXT(+)SOX2(+) neuromesodermal progenitors, PAX6(+)nestin(+) neural progenitor populations, and MEOX1(+) paraxial mesoderm populations. Wnt agonism stimulated singular axial extensions in a dose-dependent manner, and elongated organoids displayed regionalized rostral-caudal HOX gene expression, with spatially distinct hindbrain (HOXB1) expression from brachial (HOXC6) and thoracic (HOXB9) regions. CRISPR-interference-mediated silencing of the TBXT, a downstream Wnt target, increased neuroepithelial compartmentalization and resulted in multiple extensions per aggregate. Further, knock-down of BMP inhibitors, Noggin and Chordin, induced elongation phenotypes that mimicked murine knockout models. These results indicate the potent morphogenic capacity of caudalized hPSC organoids to undergo axial elongation in a manner that can be used to dissect the cellular organization and patterning decisions that dictate early nervous system development in humans.

## Introduction

Development of multicellular organisms depends on the specialization and regionalization of multiple cell types to form appropriate tissue structures and organs. In human development, this process is intertwined with the initial establishment of the anterior-posterior embryonic axis^1^, which is required for neural tube and subsequent spinal cord development. Defects in early spinal cord morphogenesis result in severe congenital abnormalities, such as spina bifida, highlighting the clinical importance of understanding human axial elongation and neural tube development. However, despite the advances gleaned from vertebrate models such as mouse and chicken, many of the molecular mechanisms and cellular behaviors that regulate spinal cord development remain unknown due to the difficulty of studying these dynamic processes following implantation^2,3^.

Experiments in model organisms have revealed that the posterior neural tube is partially generated from a unique pool of axial stem cells called neuromesodermal progenitors (NMPs). NMPs are bipotent progenitors that reside at the node-streak border within the caudal lateral epiblast and later in the chordoneural hinge of the tail bud^2,4–6^. The lack of a model for posterior neural tube development has prevented understanding specifically how NMPs regulate and coordinate the emergence of the posterior spinal cord in humans. However, studies in model systems and 2D differentiation models implicate WNT and FGF as critical factors regulating this process^7–10^.

Discoveries from embryonic studies have enabled the robust production of neuronal subtypes *in vitro*^11,12^. In parallel, organoids capable of recapitulating structures reminiscent of early embryonic tissues have been developed as high-throughput platforms for uncovering developmental patterns^13^. More specifically, protocols have capitalized on NMP differentiation^14^ to generate neuromuscular junctions^15^, mimic spinal cord dorsal-ventral patterning^16,17^, or recapitulate the complex spatiotemporal expression profiles of gastrulation (termed “gastruloids”)^18–21^. However, organoids do not perfectly recapitulate the spatiotemporal dynamics of gene expression and differentiation that occurs within embryos. Moreover, reproducibility within differentiations, between cell lines and experiments, and across labs remains a significant technical hurdle^22^.

Building on the foundation of the existing ‘gastruloid’ models currently available, we have generated an organoid model that reproduces many of the endogenous cell behaviors that generate axial elongation during human spinal cord development using both human embryonic stem cells (hESCs) and human induced pluripotent stem cells (hiPSCs). This model demonstrates extensive self-driven unidirectional growth, cell subtype specification that recapitulates NMP differentiation and neural tube morphogenesis, and regionalized gene expression profiles with distinct axial identities. This robust model of neural tube morphogenesis and axial elongation enables direct examination of features of early human spinal cord development and patterning that have to this point remained intractable.

## Results

### Wnt Agonism Induces Emergence of Axial Extension of Neuronal Organoids

Caudalization of the spinal cord and generation of NMPs are critically dependent on Wnt signaling^2,23,24^, thus we assessed whether pairing increased canonical Wnt signaling with a previously-described hindbrain differentiation protocol^11^ was sufficient to produce spinal cord cell populations. To this end, we pretreated human induced pluripotent stem cells (hiPSCs) for 48 hours with the Wnt small-molecule agonist CHIR99021 (CHIR; 4 μM) before organoid formation **(Fig. 1A,B)**. Organoids were generated by aggregating singly-dissociated cells in non-adherent pyramidal inverted wells (3,000 cells/well) followed by rotary suspension culture in 6-well plates^25^. Aggregation was considered day 0 of differentiation **(Fig. 1A)**. After 5 days in suspension culture, pronounced singular extensions emerged from the CHIR-pretreated organoids **(Fig. 1C)**. Elongated organoids were continuous, with linear striations extending longitudinally from a radially symmetric anterior aggregate. Histological analysis revealed multiple internal elongated epithelial compartments separated by regions devoid of cells **(Fig. 1C)**. To ensure elongation was not due to organoid fusion in bulk culture, organoids were imaged continuously over 48 hours which revealed robust elongation in 83% of imaged wells as measured by axis ratios above 1.6 at the end of the time-course **(Fig. 1D, Fig. S1A-B, Movie S1-2)**. Because NMPs can be generated *in vitro* in as little as 48 hours^12^, we examined whether CHIR treatment generated a detectable population of NMPs immediately following aggregation. Indeed, SOX2(+)TBXT(+) NMPs were present in CHIR organoids at day 0, whereas in the absence of CHIR treatment, organoids lacked NMPs and failed extend **(Fig. S2A,B)**. To ascertain whether extensions resulted from polarized populations of proliferating cells, we examined EdU incorporation and phospho-histone 3 presence on days 6, 7 and 9 of differentiation **(Fig. S3A)**. A two-hour EdU pulse labeled >50% of cells in extending organoids, where proliferation was observed increasingly at the outer edges of day 6 and 7 extending aggregates (quadratic model fitting, p < 0.001) where there is roughly a 15% increase in EdU labeling from center to edge of the organoids. Additionally, this was recapitulated to a lesser degree (1% or less change, quadratic model fitting, p = 0.015) when examining actively dividing cells via phospho-histone 3 presence. Furthermore, extensions occurred in organoids of various sizes **(Fig. S4B)** and extended more robustly when cultured at low density **(Fig. S4A)**, suggesting that density-mediated signaling parameters, such as paracrine effects or nutrient availability, likely induce and maintain axial extensions, similar to the documented role of glycolysis in vertebrate axial elongation^26^. Overall, we generated an organoid model that robustly extends in a variety of culture conditions due to the early presence and number of SOX2(+)TBXT(+) NMPs that seem to account for the generation of the elongation phenotype.

**Figure 1:**
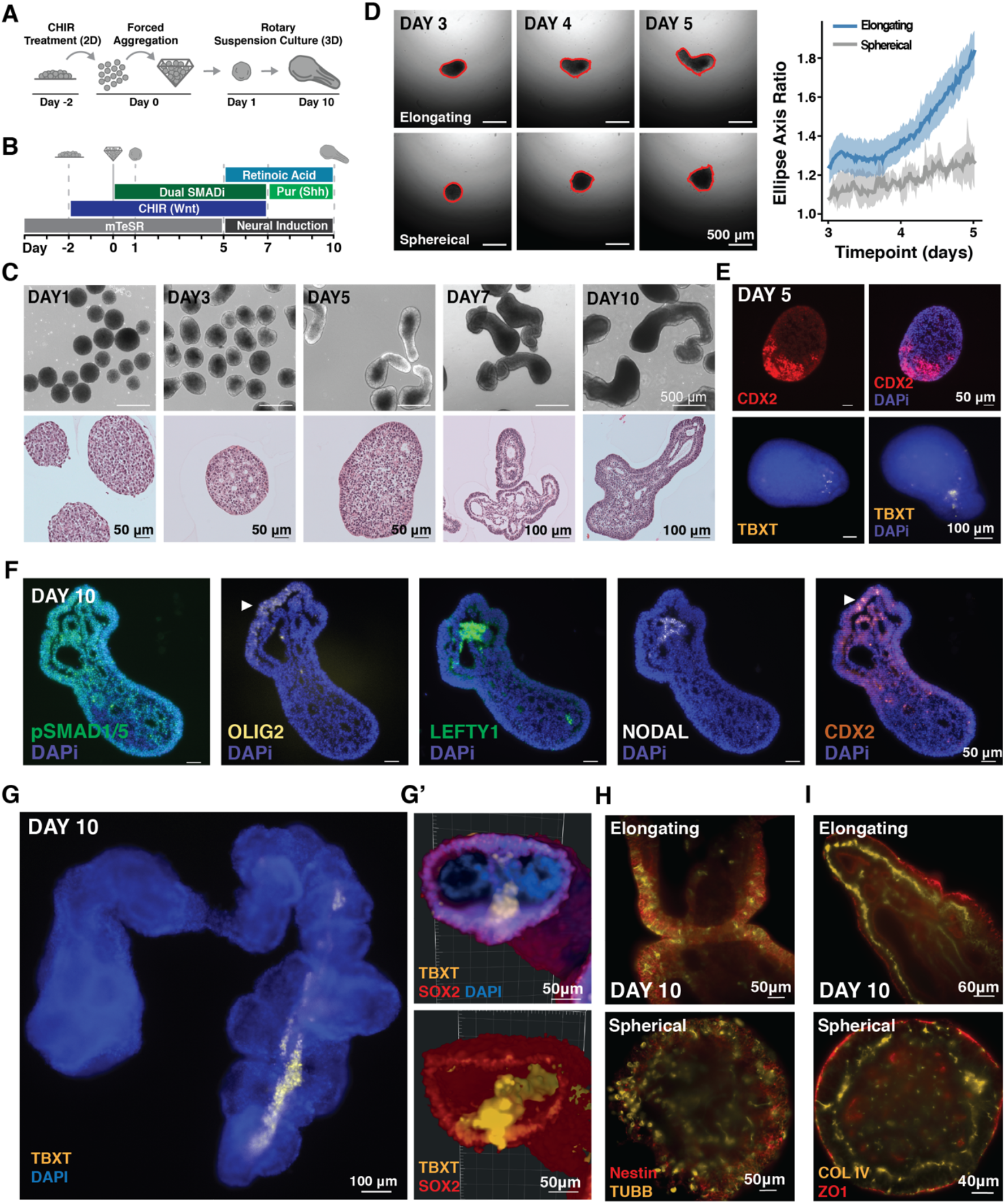
**A-B)** Schematic of experimental set up and differentiation protocol. **C)** Brightfield and histological images of extending organoids. **D)** (LEFT) Frames from video time-course tracking organoid extensions in static 96 well plate culture where red outline defines measurements taken. (RIGHT) Quantification of ellipse axis ratios from day 5 to 7 of extending (blue) and non-extending (grey) organoids. Solid lines represent the mean ratio (dark color) with 95 percent confidence interval (light color); n= 2 non-elongating, n= 15 elongating. **E) I** mmunofluorescence images at day 5 of differentiation from paraffin sections in extending organoids examining CDX2 and TBXT localization. **F) I** mmunofluorescence images at day 10 of differentiation from paraffin sections in extending organoids examining axis and developmental patterning markers. **G)** Fluorescence image of TBXT streak traveling down the length of organoid **G’)** 3D reconstruction cross section of the posterior of extending organoids based on light-sheet microscopy displaying TBXT+ streak. **H-I)** Optical sections from light-sheet microscopy of elongating (top) and spherical (bottom) organoids stained for Nestin and bIII-Tubulin (TUBB) or COL IV and ZO1.

### Organoids Display Markers of Tissue and Cellular Polarity

Based on the robust axial elongation, we next examined whether extending organoids generated an anterior-posterior (A-P) axis analogous to embryonic development. The posteriorly expressed gene CDX2^27^ was detected in a polarized manner from day 5 to day 10 of differentiation **(Fig. 1E,F)**. Furthermore, within extending organoids, clusters of NODAL- and LEFTY-positive cells were observed at day 10, suggesting the presence of an organizer-like population **(Fig. 1F, S5B)**. In fact, multiple clusters of NODAL-positive cells were present in organoids with multiple extensions **(Fig. S5A)**. In day 10 extending organoids, compartments of cells expressing the ventral spinal cord marker, OLIG2, lined the epithelial cysts, and a polarization of pSMAD1/5 expression was detected along the length of the organoid **(Fig. 1F)**. These observations suggest that the extending organoids exhibit morphogenic characteristics of the emergent cell populations that establish the embryo’s anterior-posterior axis.

To interrogate how differences in the identity and spatial distribution of NMP populations in extending organoids contributed to differential organoid morphologies, we imaged extending and non-extending organoids at day 7 of differentiation **(Fig. 1G-G’, S5C)**. In most extending organoids, the TBXT(+) population formed a streak along the protruding extension and accumulated at the posterior end of the extension, reminiscent of the notochord and previous gastruloid reports^18^. Additionally, regions of SOX2(+)TBXT(+) NMPs were found to be immediately adjacent to the TBXT(+) streak **(Fig. 1G’)**. In contrast, TBXT(+) cells in non-extending organoids were sparsely distributed across the surface of the spherical organoid with no clearly identifiable regions of SOX2(+)TBXT(+) co-expression **(Fig. S5C)**. After 10 days, extending and non-extending organoids both contained cells expressing the neuronal progenitor markers Nestin and beta 3-tubulin, indicating that the majority of cells differentiated to a neural fate **(Fig. 1H, S6B, Movies S5-6)**. However, in the extending organoids, the neural-committed cells organized into distinct compartmentalized layers surrounding lumens within the tissue-level organoid extensions.

We examined apical-basal cell polarity by staining for collagen IV (COL IV), a marker of the basement membrane, and zona occluden-1 (ZO-1), a marker of tight junctions at cellular apical domains **(Fig. 1I, Fig. S6A)**. An increase in basement membrane deposition and cell polarization was observed between days 7 and 10 of differentiation in extending organoids **(Fig. S6C)**. After 10 days, ZO-1 was apparent at the apical domain of cells in the outer epithelial layers of the extensions, as well as in cells facing the internal lumens. Distinct layers of basement membrane marked by collagen IV were observed basally to ZO1, forming a bilayer between lumens and organoid exterior **(Fig. 1I, Fig. S6A,C, Movies S7-8)**. Bilayers of cells were also apparent in non-extending organoids, but to a lesser extent since the internal lumens were much smaller. Overall, these patterns indicate that extending organoids undergo tissue polarization via axial extensions that consist of polarized sheets of neuroepithelial cells that generate organized basement lamina.

### Organoids Display Regionalized HOX Patterning

To examine transcriptional differences between the two poles of extending organoids, we manually dissected the “anterior”, or main organoid body, from the “posterior”, or extending portion, of day 9 organoids **(Fig. S7A,B)** and performed comparative qPCR. Of the genes tested, only HOXA11, a marker of the lumbar region, displayed a significant difference in expression between the anterior and posterior fragments, likely because of variability between individual organoids **(Fig. S7C)**. To gain a more nuanced understanding of HOX expression in extending organoids, we examined transcripts marking hindbrain (HOXB1), brachial (HOXC6), and thoracic (HOXB9) regions of the neural tube via RNAscope at days 7 and 10 **(Fig. 2A, S8A)**. While non-extending organoids expressed both hindbrain and brachial HOX genes, their expression was distributed radially, and the expression of HOXB9 was delayed relative to extending organoids **(Fig. S8B)**. In contrast, HOXB1 (hindbrain) was enriched in the central mass of extending organoids, and HOXC6 (brachial) and HOXB9 (thoracic) were enriched in the extensions **(Fig. 2A)**. Interestingly, HOXC6 and HOXB9 often overlapped at day 7 within extending organoids. However, by day 10 HOXB9 was more pronounced than HOXC6 in the extension regions, suggesting a transition to a more posterior fate. These data indicate that while organoid extension is not a requirement for posterior HOX expression, extension enables the stratification of distinct HOX domains.

**Figure 2:**
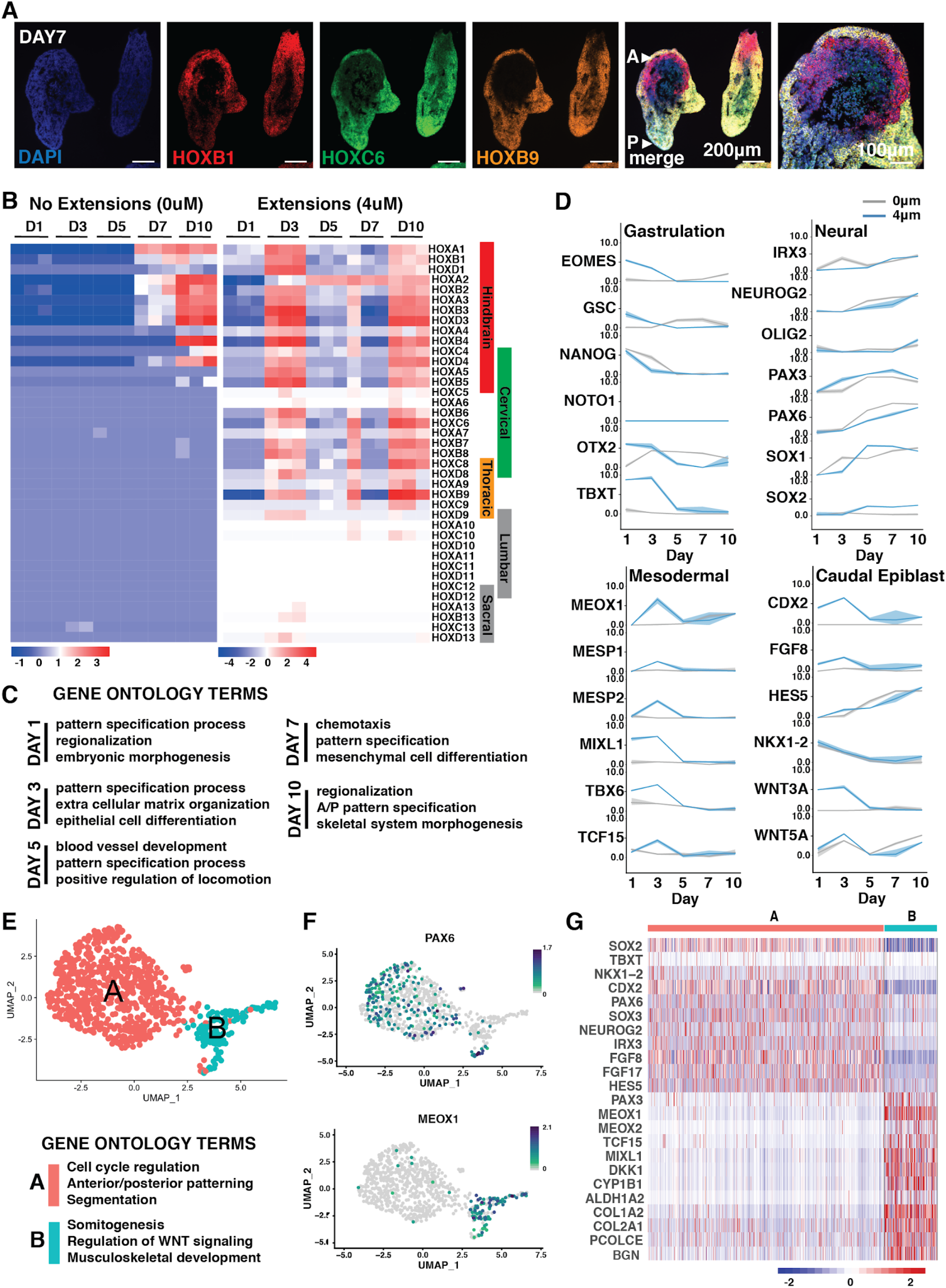
Gene expression differences within extending organoids. **A)** RNAscope of sectioned organoids cultured at low density with probes for HOX genes marking different regions of the spine at day 7 of differentiation (arrows mark anterior (A) and posterior (P) axis of organoids). **B)** Heatmap of HOX gene expression in organoids with and without CHIR treatment from bulk RNAseq **C)** Line graphs of gene expression change in markers associated with Gastrulation, Neural, Mesodermal, and Caudal Epiblast fates (grey line indicates 0uM CHIR and blue line indicates 4uM CHIR; solid line indicates mean, shading represents 95% confidence bounds; axis displayed are log2 counts). **D)** Significant (p < 0.05) Gene Ontology classifications derived from the most differentially upregulated genes in extending organoids from bulk RNAseq. **E)** UMAP of n=789 cells from extending organoids exposed to a lower concentration of CHIR (2μM) at day 10 of differentiation from single-cell RNAseq. Gene Ontology terms assigned to identified clusters. **F)** UMAPs showing cells expressing PAX6 and MEOX1. Color scale indicates a normalized increase in log2 fold change from min expression to max expression of the respective gene. **G)** Heat map showing normalized expression levels of genes associated with spinal neuron specification, caudalizing factors, and mesoderm specification from single-cell RNAseq.

### Developmental Timing and Maturation of Organoids

To obtain a more detailed view of HOX gene expression over time, we performed bulk RNA sequencing, comparing organoids with and without CHIR treatment. As observed in previous imaging, timing and extent of HOX expression varied between the two conditions. Non-extending organoids began to express detectable HOX transcripts at day 7 and adopted a hindbrain identity upon the experiment’s conclusion at day 10 **(Fig. 2B)**. In contrast, extending organoids expressed both hindbrain and cervical HOX transcripts as early as day 3, with a notable upregulation in thoracic HOX genes by day 10. This temporal change in HOX emergence, in conjunction with the RNA-scope data, suggests that the spatiotemporal onset of HOX expression in extending organoids recapitulates the progression of anterior-to-posterior elongation of *in vivo* development.

Analysis of the top divergent genes between the extending and non-extending organoids **(Fig. S9A)** revealed unique components of pattern specification in the extending condition, as indicated by Gene Ontology **(Fig. 2C)**. Day 1 extending organoids were enriched for genes involved in pattern specification and embryonic morphogenesis, and by day 10 included hallmarks of anterior-posterior pattern specification. Extending organoids upregulated WNT3 and WNT5B signaling at day 1, whereas the non-extending organoids upregulated WNT5B at day 5 **(Fig. 2D, S9B)**. Extending aggregates were enriched for the caudal marker CDX2, while non-extending organoids were enriched for the forebrain marker OTX2 at later timepoints **(Fig. 2D, S9A)**. Further, elongating organoids passed through a goosecoid (GSC) and TBXT-high stage at day 1 and more specific lineage markers arose at day 3. Finally, neural markers (IRX3, NEUROG2, OLIG2, PAX3, PAX6, SOX1, SOX2) remained high as the differentiation progressed. Together, this analysis indicates that early upregulation of Wnt signaling induces pattern specification in organoids that mirrors many aspects of *in vivo* morphogenesis and favors caudal neurogenesis.

### Extending Organoids Consist Exclusively of Neural and Mesodermal Lineages

To examine cell diversity in extended organoids, we performed single-cell transcriptomics on extending day 10 organoids. The single cell transcriptomic data recapitulated HOX gene expression profiles previously observed within extending organoids **(Fig. S10A)**. Shared nearest neighbor computation revealed two clusters **(Fig. 2E)**; one (cluster A) largely distinguished by cell cycle regulation and the other (cluster B) by somitogenesis, based upon Gene Ontology terms. Neural markers, such as SOX2, were expressed in both clusters **(Fig. S10B,C)**, but, cluster A had higher expression levels of genes associated with neural tube fates (SOX2, IRX3, PAX6, NEUROG2) and the caudal lateral epiblast (CDX2, NKX1.2 and FGF8) **(Fig. 2F,G)**. In contrast, MIXL1, MEOX1, CYP1B1, MEOX2, and PAX3 were almost exclusively expressed in cluster B, indicating that this cluster contained mesodermal cells **(Fig. 2F,G; S10B,C)**. We did not observe expression of endoderm-specific markers such as SOX17, and only very low expression of FOXA1 and PAX9 in <1% of cells **(Fig. S10C)**, indicating that, in contrast to previously reported gastruloid models^18–20^, these extending organoids consisted entirely of neural and mesodermal cell types. Overall, these data indicate that extending organoids transcriptionally mimic the caudal epiblast and contain cell populations that appear to be exclusively associated with both mesoderm and neural tube, similar to the embryo during axial extension.

### Wnt Signaling Influences Extension and Axis Identity Across Stem Cell Lines

To determine the robustness of WNT signaling effects on axial elongation, we differentiated two iPSC lines (WTC and WTB) and two hESC lines (H1 and H7) into neural organoids in the presence of varying amounts of CHIR. The threshold of CHIR required to reliably generate extensions varied between lines **(Fig. 3A)**, highlighting cell-line-specific responsivity to Wnt agonism. Whereas the WTC line produced elongations at a 2-4μM dose of CHIR, all other lines tested only began to elongate when the CHIR dose was increased to 6μM. Further, increasing CHIR dosage increased both the degree of extension **(Fig. 3B,C’,D’)** and the frequency of SOX2(+)TBXT(+) cells at the start of the differentiation **(Fig. 3C,D)**. Interestingly, the WTC line demonstrated a maximum threshold for CHIR-induced extension, as 6μM CHIR treatment reduced extensions **(Fig. 3A,C)**. Importantly, all cell lines displayed polarized organization with internal epithelialization and cavitation at their respective condition that permitted extension **(Fig S11A)**. To further characterize the polarized extension across cell lines, we found that the H1 ESC line displayed similar regionalized expression of axial markers as seen in the WTC iPSC line **(Fig. S11B)**.

**Figure 3:**
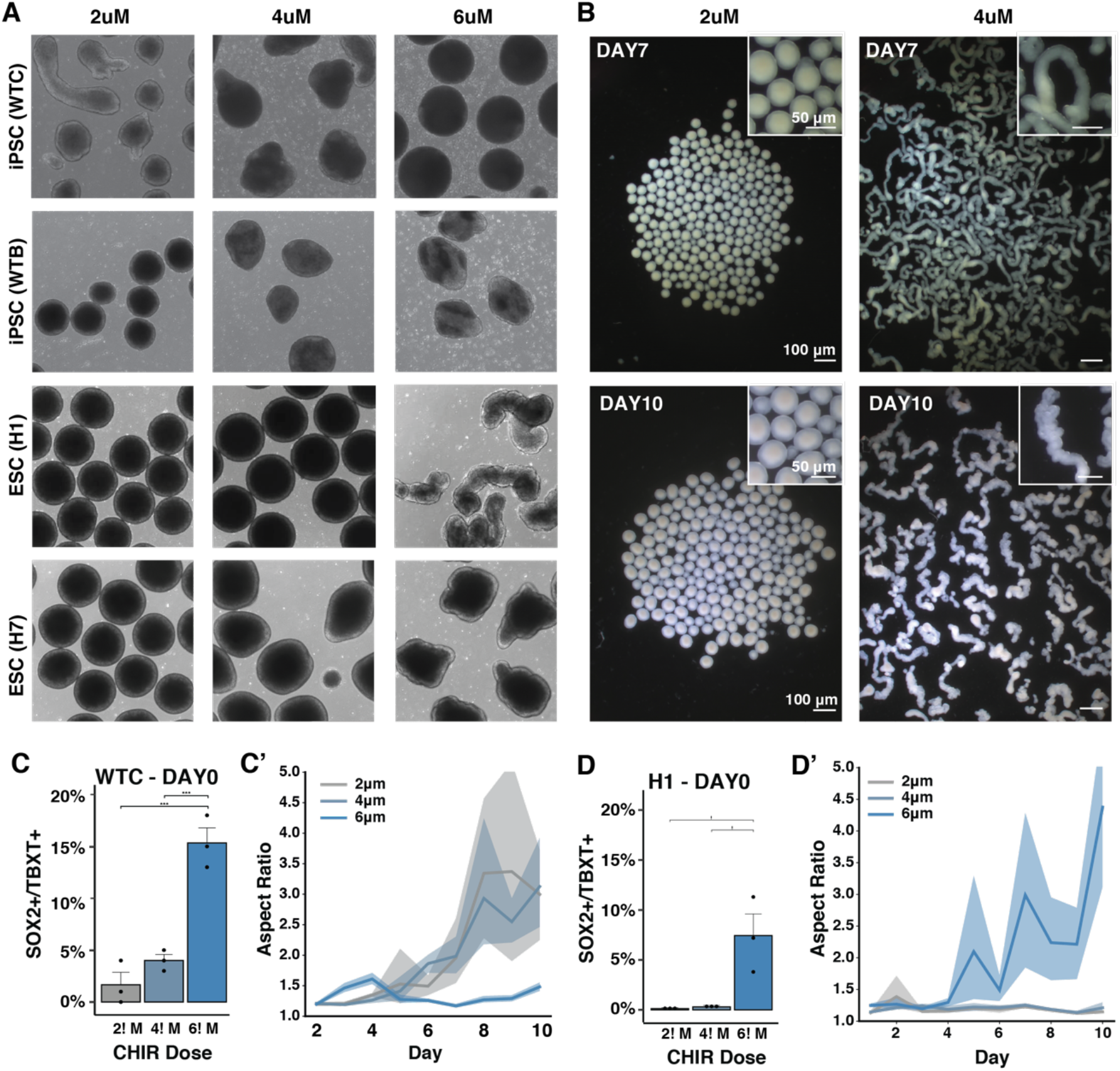
Wnt mediated increase in extensions across stem cell lines. **A)** Brightfield images at day 7 of differentiation with increased CHIR doses conducted in the hiPSC and ESC lines. **B)** Stereoscope images of extending and non extending organoids in two different CHIR doses. **C)** Quantification of SOX2(+)TBXT(+) cells at increasing CHIR doses by FLOW cytometry in the WTC hiPSC or H1 ESC line at day 0. **D)** Quantification of the length of extensions in WTC or H1 organoids shown as radius ratio (ie. major to minor) with increasing doses of CHIR. (Solid line indicates mean, shading represents 95% confidence bounds).

To examine differences in expression across Wnt conditions, we stained sections of WTC organoids at day 10 treated with 2μM, 4μM or 6μM of CHIR **(Fig. S12)**. Organoids exposed to 4μM and 6μM CHIR expressed SOX2 and PAX6 in distinct regions of the organoids, whereas organoids exposed to 2μM CHIR displayed SOX2 and PAX6 expression throughout, suggesting a non-neuronal population emerging in higher Wnt signaling environments. All organoids displayed N-cadherin and beta III-tubulin expression, indicating the emergence of maturing populations of neurons; however, regions without N-cadherin or beta III-tubulin were more prevalent in organoids exposed to higher doses of CHIR, suggesting a Wnt-dependent reduction in the overall neural population. Altogether, the level of Wnt pathway activation that allows for robust emergence of an NMP progenitor pool and organoid extension varies across both hiPSC and ESC lines - a common phenomena for most differentiation protocols^22^.

### Wnt and BMP Signaling Influences Axis Identity and Degree of Extension

As the body axis extends, gradients of key signaling molecules are established across the embryo. Initially, Wnt signaling is activated throughout the epiblast by extraembryonic BMP and later localized to the posterior embryo where Wnt directly upregulates the expression of TBXT, which in turn upregulates both Wnt and FGF, reinforcing and maintaining their posterior to anterior gradients^28^. The importance of these signaling relationships for proper axial elongation are demonstrated in mouse knockout models. TBXT knockout mice have severely disrupted trunk morphogenesis, failing to generate the notochord and posterior somites^29–33^. When the dorsal inhibitor of BMP, Noggin, is knocked out, murine models have increased posterior BMP signaling, failure of neural tube closure, loss of robust dorsal-ventral patterning, and elongation of the developing tail^9^. Deletion of the BMP shuttling molecule, Chordin, results in a shortened body axis, a ventralized phenotype, and underdeveloped anterior spine^34^. Thus, to interrogate the role of these signaling pathways in a human system, we independently knocked down TBXT, Chordin, or Noggin via an inducible CRISPR interference system in human iPSCs^35–37^.

First, we integrated RNA guides targeting TBXT into a Lamin-B GFP-labeled WTC hiPSC line harboring a doxycycline (DOX) inducible CRISPR interference system (CRISPRi)^35,36^. Media supplemented with DOX for five days prior to cell aggregation reduced the number of TBXT(+) cells by 90% **(Fig. 4A,B)**, and the frequency of TBXT(+)SOX2(+) cells was also significantly reduced **(Fig 4C)**. Interestingly, the TBXT knockdown organoids displayed multiple extensions emerging at day 5 **(Fig 4D)**. These multi-extension aggregates maintained a reduced level of TBXT expression through day 10 and had large cavities devoid of cells **(Fig. 4E,F)**. At day 7, the wildtype control demonstrated pockets of expression of the axis organizer LEFTY, whereas the TBXT knockdown aggregates displayed reduced expression of LEFTY that was distributed throughout the organoids **(Fig. 4G)**. Isogenic control and TBXT knockdown organoids both expressed the neuronal marker Nestin, the caudal marker CDX2, and basement membrane marker fibronectin **(Fig 4H)**. However, there was a reduction in caudal HOX genes in the knockdown organoids at days 5 and 10 of differentiation, as well as a reduction in CDX2 and FGF8 expression **(Fig. 4I, Fig. S13)**, reminiscent of the absent posterior structures in TBXT knockout mice. Interestingly, organoids maintained expression of members of the Wnt pathway, such as AXIN2 and WNT3, suggesting that TBXT is necessary in axial organoids for unidirectional extension and caudal fate specification, but not for maintaining Wnt expression and signaling.

**Figure 4:**
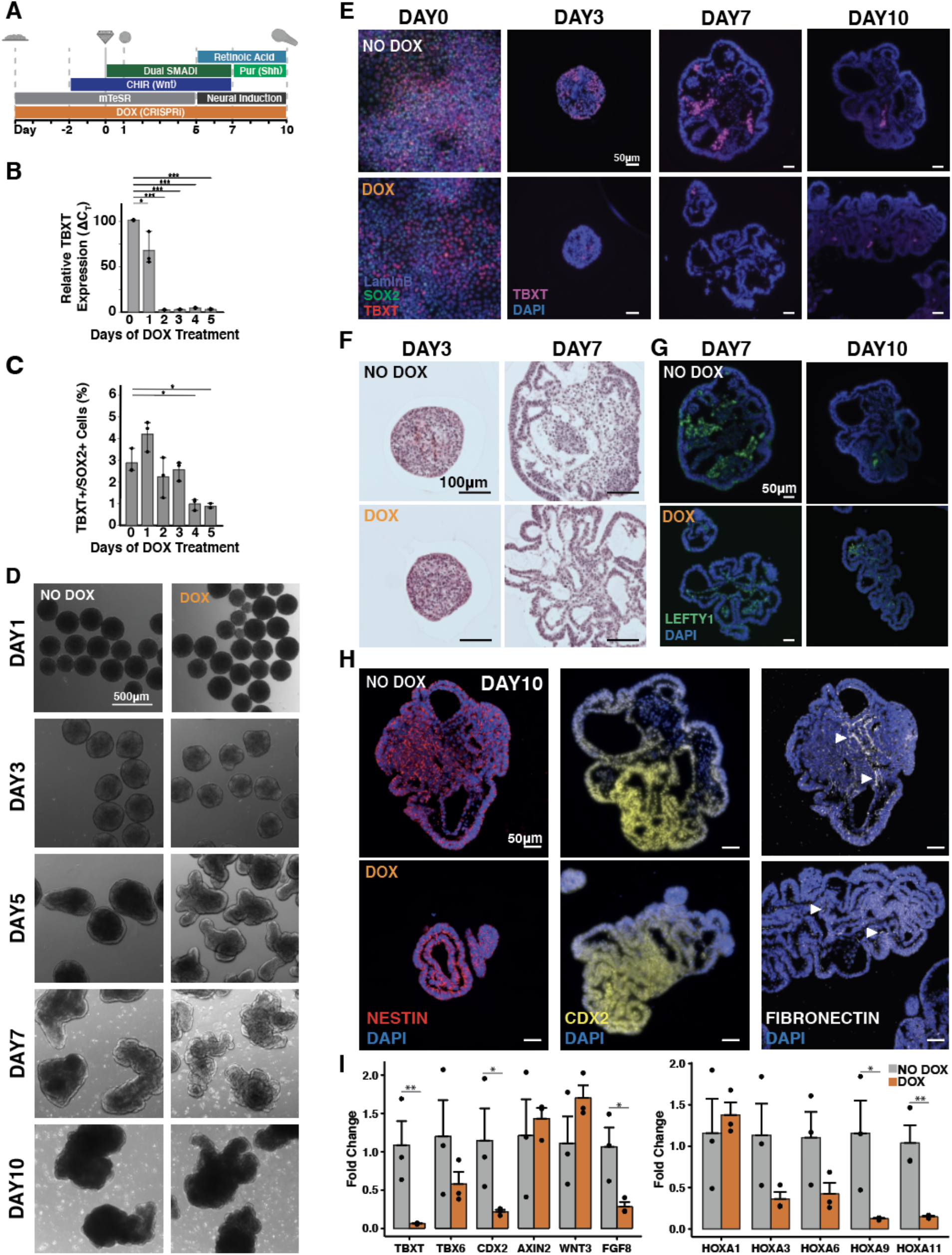
TBXT knockdown leads to multiple elongations. **A)** Schematic of differentiation protocol timeline with knockdown induction. **B)** Quantification of TBXT knockdown efficiency by reduction in mRNA expression levels measured by qPCR, each collected on day 0 of differentiation. **C)** Quantification by FLOW of TBXT(+)SOX2(+) cells with DOX treatment over time, each collected on day 0 of differentiation. **D)** Brightfield images of differentiation time course of knockdown hiPSC lines. E) Immunofluorescence staining of fixed cells (Day 0) or paraffin sections (Days 3-10) for TBXT over time in wildtype and KD organoids. F) Histological sections of wildtype and KD organoids at day 3 and 7 of differentiation. G) Immunofluorescence staining of paraffin sections for LEFTY at day 7 and 10 in wildtype and KD organoids. H) Immunofluorescence staining of paraffin sections for NESTIN, CDX2, and Fibronectin at day 10 in wildtype and KD organoids. I) qPCR quantification of HOX genes and genes related to axial pattern specification from day 10 organoids with and without DOX treatment (* signifies p-value < 0.05, ** signifies p-value < 0.005, *** signifies p-value < 0.0005).

To interrogate the BMP pathway, we knocked down Noggin and Chordin separately **(Fig. S14B)**. Both Chordin and Noggin knockdown resulted in increased extension compared to wildtype organoids **(Fig. S14C)**. To further examine extension dynamics, we allowed the organoids to continue to grow for an additional 7 days, to a total of 17 days of differentiation. Noggin knockdown organoids extended continuously, becoming more polarized through day 13 relative to isogenic control organoids. The pronounced extensions in Noggin knockdown organoids appeared analogous to the elongated tails of Noggin knockout mice. In contrast, Chordin knockdown organoids initially rapidly developed outgrowths without clear anterior-posterior axis identity, but then halted elongation after day 10 of differentiation, a morphogenic response reminiscent of the shortened body axis of Chordin knockout mice **(Fig. S14D)**.

Because of the role of BMP signaling in generating the dorsal-ventral axis of the spinal cord, progenitor domain gene expression was assessed by qPCR. At day 17, expression of LHX2, a marker of dI1 dorsal interneurons, did not significantly change in either knockdown conditions. However, expression of LHX5, a marker of V0 ventral interneuron populations, significantly increased in both Noggin and Chordin knockdown organoids **(Fig. S14E,F)**. Although markers and regulators of neural maturation, Sox2, Pax6, and Hes1, were not significantly upregulated in the knockdowns, LHX5 and EN1, transcription factors that mark the central V0 and V1 domains, were upregulated. Interestingly, genes associated with the more extreme poles of the dorsal-ventral axis (LHX2, CHX10, and OLIG2) did not show significant differences, suggesting a potential change in the threshold of BMP signaling that affects more central regions within the dorsal-ventral axis. Altogether, these results suggest that BMP signaling impacts both the anterior-posterior and dorsal-ventral progenitor emergence of the extending human organoids in a manner that recapitulates cellular patterning of murine neural tube development.

## Discussion

Here, we demonstrate that hiPSC-derived organoids can model axial extension via the specification of progenitors that organize axial elongation and neural tube formation. This model enables robust axial extension as a function of both culture density and Wnt signaling. Within extending organoids, we observed a persistent TBXT(+)SOX2(+) NMP population, reminiscent of populations present in the caudal lateral epiblast and in the tail bud during axial elongation. Extending organoids displayed regionalized co-linear patterning and temporal expression of HOX genes. Single-cell RNA sequencing confirmed the presence of caudal transcriptomic profiles and populations of neuronal progenitors and paraxial mesoderm. Finally, we demonstrated that the axial extension of organoids is influenced by Wnt and BMP signaling, such that perturbation of endogenous TBXT, Chordin, or Noggin influences relative expression of neuronal markers along the anterior-posterior and dorsal-ventral axis, as observed in parallel mouse models^9,30–34^. Overall, the spinal organoids described herein provide reliable and faithful models of the gene regulatory networks and multicellular structural organization that contribute to the development of the primitive human central nervous system.

Stem cell-derived organoid models that reflect multiple organ systems^15,38–40^ and stages of development^17–20,41^ can now be used to investigate aspects of human embryogenesis that were previously elusive due to poor tissue quality, technical difficulties isolating and imaging embryonic tissues, and ethical concerns regarding tissue procurement. Organoid platforms are amenable to varying culture conditions, genetic modification, or small molecule drug screens, enabling the systematic manipulation of experimental conditions at higher scale and speed than possible in most model organisms. Our work demonstrates that as multiple examples of axial elongation are developed^17–20,42,43^, refining the protocols of organoid culture and differentiation can lead to more complex models than previously possible, and pushes the limits of organoid utility beyond cell signaling toward coordinated multicellular organization and morphogenesis en route to structurally and functionally mature tissues.

## Methods

### Human Induced Pluripotent Stem Cell Line Generation and Culture

All work with human induced pluripotent stem cells (iPSCs) or human embryonic stem cells (ESCs) was approved by the University of California, San Francisco Human Gamete, Embryo, and Stem Cell Research (GESCR) Committee. Human iPSC lines were derived from the WTC11 line (Coriell Cat. #GM25256), the WTB line (Conklin Lab)^44^, and the Allen Institute WTC11-LaminB cell line (AICS-0013 cl.210) and the human ESCs H7 and H1 (WiCell, Madison, WI). All cell lines were karyotyped by Cell Line Genetics and reported to be karyotypically normal. Additionally, all cell lines tested negative for mycoplasma using a MycoAlert Mycoplasma Detection Kit (Lonza).

Human iPSCs were cultured on growth factor reduced Matrigel (Corning Life Sciences) and fed daily with mTeSRTM-1 medium (STEMCELL Technologies)^45^. Cells were passaged by dissociation with Accutase (STEM CELL Technologies) and re-seeded in mTeSRTM-1 medium supplemented with the small molecule Rho-associated coiled-coil kinase (ROCK) inhibitor Y-276932 (10 μM; Selleckchem)^45^ at a seeding density of 12,000 cell per cm^2^.

The generation of the TBXT, Chordin and Noggin CRISPRi lines first involved TALEN mediated insertion of the CRIPSRi cassette pAAVS1-NDi-CRISPRi (Gen2) (Addgene) to the AAVS1 locus of the Allen Institute WTC11-LaminB cell line. Following antibiotic selection of clones that received the CRIPSRi cassette, CRISPRi gRNAs were generated targeting Noggin and Chordin (Table S2) using the Broad Institute GPP Web Portal and cloned into the gRNA-CKB (Addgene) following the previously described protocol^36^. Guide RNA vectors were nucleofected into the LaminB CRISPRi iPSC line using a P3 Primary Cell 96-well NucleofectorTM Kit (Lonza) and the 4D Nucleofector X Unit (Lonza) following manufacturer’s instructions. Nucleofected cells were allowed to recover in mTeSRTM-1 medium supplemented with Y-276932 (10 μM) and then underwent antibiotic selection with blasticidin (ThermoFisher Scientific; 10 μg/ml) following the previously published protocol^35,36^. Knockdown efficiency was evaluated by addition of doxycycline to the daily feeding media over the course of 5 days, collection of mRNA, and subsequent quantification of gene expression by qPCR.

### Organoid Differentiation

Organoid differentiations were a modified protocol of a previously published spinal cord interneuron differentiation protocol^11^. Human iPSCs were seeded at 125000 cells/cm^2^ in mTeSRTM-1 medium supplemented with the small molecule Rho-associated coiled-coil kinase (ROCK) inhibitor Y-276932 (10 μM; Selleckchem) and small molecule GSK inhibitor CHIR99021 (2μM, 4μM, or 6μM; Selleckchem). Two days later, cells were singularized with Accutase (STEMCELL Technologies), counted using a Countess II FL (Life Technologies), and seeded into 800μm X 800μm PDMS microwell inserts in a 24 well plate (~270 wells/insert)^25^. After ~18 hours, condensed organoids were transferred to rotary culture in 6-well plates in mTeSRTM-1 medium supplemented with Y-276932 (10 μM; Selleckchem), CHIR99021 (2μM, 4μM, or 6μM; Selleckchem), ALK5 small molecule inhibitor SB431542 (10μM, Selleckchem), and small molecule BMP inhibitor LDN193189 (0.2μM, Selleckchem) at an approximate density of 270 aggregates per well unless otherwise mentioned in figure legend. Organoids were fed every other day for up to 17 days. Y-276932 was removed from the media at day 3. At day 5 organoids were transferred to Neural Induction Media (DMEM F:12 (Corning), N2 supplement (Life Technologies), L-Glutamine (VWR), 2μg/ml heparin (Sigma Aldrich), non-essential amino acids (Mediatech INC), penicillin-streptomycin (VWR), supplemented with fresh 0.4μg/ml ascorbic acid (Sigma Aldrich) and 10ng/ml brain derived neurotrophin factor (BDNF, R&D Systems)) supplemented with CHIR99021 (2μM, 4μM, or 6μM; Selleckchem), SB431542 (10μM, Selleckchem), and LDN193189 (0.2μM, Selleckchem). From day 7 onwards, organoids were fed with Neural Induction Media supplemented with retinoic acid (10nM, Sigma Aldrich), purmorphamine (300nM, EMD Millipore) and N-[N-(3,5-difluorophenacetyl)-L-alanyl]-S-phenylglycine t-butyl ester (DAPT D5942, 1μM, Sigma-Aldrich).

### Organoid Elongation Imaging and Quantification

Day 5 organoids from a single 10cm dish were individually transferred using wide bore pipette tips into the center 60 wells of an uncoated ultra-low attachment 96-well plate (Corning), seeding exactly one organoid per well, with the remaining organoids maintained in rotary culture through day 7. Using an inverted Axio Observer Z1 (Zeiss) microscope with incubation (Zeiss Heating Unit XL S, maintained at 37°C, 5% CO_2_), all 60 wells were imaged using an AxioCam MRm (Zeiss) digital CMOS camera at 5x magnification (NA 0.16, 2.6 μm x 2.6 μm per pixel). Each well was imaged in TL Brightfield every 20 minutes for 48 hours giving a total of 145 frames. At the end of imaging (day 7), 31 organoids from the parallel rotary culture were imaged at 5x to generate a comparison image set.

To segment the organoids, all well images were first aligned by fitting a truncated quadratic curve to the average image intensity, then solving for the peak of maximum intensity, which was assumed to be the well center. Next, the average lighting inhomogeneity was calculated as the pixel-wise median of all 60 aligned well images, which was then subtracted from the individual aligned frames. After background correction, individual organoids were isolated by finding objects brighter than 0.83% of maximum intensity, but less than 3.0% of maximum intensity, with object size greater than 2,000 pixels, eccentricity greater than 0.1, and solidity greater than 40%. A bounding box 2 mm x 2 mm around the center of each of these objects was calculated and all frames of the time series cropped to this bounding box to reduce memory usage. To detect the region of maximum motion in the time series, the difference image between each pair of sequential images was calculated, and then the pixel wise standard deviation was calculated over all difference images in a given region. This standard deviation image was then thresholded at between 0.01 and 0.03 (AU) depending on the remaining lighting inhomogeneity in the image, producing a ring-shaped mask around the periphery of each organoid. Finally, using the interior of the mask as the organoid seed and the exterior as the background seed for the first frame, organoids were segmented using anisotropic diffusion^47^, evolving the foreground and background seeds using the contour calculated from the previous frame for subsequent segmentations. Segmentation, labeling, and metrology were all performed using the python package sckit-image^48^.

Segmentations were manually inspected for accuracy, with 45 of 60 determined as having no or only minor flaws, with the remaining 15 excluded from automated analysis. Using the high-quality segmentations only, each organoid time series was then analyzed to examine geometry change over time. For each contour at each time point, we calculated contour area, contour perimeter, minimum, maximum and mean distance from contour center of mass to the perimeter. As additional non-dimensional measures of shape, we calculated the ratio of maximum to minimum radius and organoid circularity. Organoids were also manually classified as “extending”, “partially extending”, or “non-extending” by examining each video. Organoids assigned to “extending” exhibited at least one, and at most two large protuberances that extended at least 100μm from the main body. Partially extending organoids exhibited at least one, and often many protuberances, all of which failed to extend robustly past the 100μm demarcation. Non-extending organoids were any organoids that failed to generate any extensions over the observation period.

### Dissection of Extended Organoids

Dissections were performed on a SteREO Discovery.V8 Manual Stereo Microscope (ZEISS) and images were taken with an EP50 Microscope Digital Camera (Olympus). Day 9 organoids were transferred into 12 mL of 37°C PBS in a 10 cm dish. 2 mL of warmed PBS were also added to 2 wells of a 6-well plate. Using two fine point forceps, polarized elongated aggregates were immobilized and pulled or pinched apart. Using a p200 pipette with a cut tip, the ‘anterior’ and ‘posterior’ halves of the organoids were collected and transferred to separate wells of the 6-well plate. After ~25 organoids were dissected, the collected halves were transferred to Eppendorf tubes and suspended inRLT buffer (RNAeasy Mini Kit; QIAGEN) for RNA extraction. Remaining, whole undissected organoids were also collected and suspended in RLT buffer for comparison. Standard RNA extraction was performed following manufacturer specifications.

### Real Time Quantitative Polymerase Chain Reaction

Total RNA was isolated from organoid samples using an RNAeasy Mini Kit (QIAGEN) according to manufacturer’s instructions. Subsequently, cDNA was generated using an iScript cDNA Synthesis kit (BIORAD) and the reaction was run on a SimpliAmp thermal cycler (Life Technologies). Quantitative PCR reaction using Fast SYBR Green Master Mix (ThermoFisher Scientific) and run on a StepOnePlus Real-Time PCR system (Applied Biosciences). Relative gene expression was determined by normalizing to the housekeeping gene 18S rRNA, using the comparative threshold (CT) method. Gene expression was displayed as fold change of each sample versus control. The primer sequences were obtained from the Harvard Primer bank or designed using the NCBI Primer-BLAST website (Table S1).

Due to low RNA abundance in the dissected organoids, PreAmp Master Mix (FLUIDIGM) was used to amplify cDNA. Briefly, 20ng of cDNA were amplified per 5μL reaction volume for 15 cycles on a SimpliAmp thermal cycler (Life Technologies). Amplified cDNA was diluted 5-fold using nuclease free water. 1μL of amplified diluted cDNA was used for each 20μL quantitative PCR reaction using Fast SYBR Green Master Mix (ThermoFisher Scientific) and run on a StepOnePlus Real-Time PCR system (Applied Biosciences), following normal qPCR methods described above.

### Histology, Immunocytochemistry and Imaging

Organoids were fixed with 4% paraformaldehyde (VWR) for 40 minutes, washed three times with PBS. Organoids to be used for histology were embedded in HistoGel Specimen Processing Gel (Thermo Fisher) prior to paraffin processing. Parafin embedded samples were sectioned in 5m sections, and subsequently stained for H&E. For immunofluorescent staining, slides were deparaffinized as for H&E staining. Epitope retrieval was performed by submersing slides in Citrate Buffer pH 6.0 (Vector Laboratories) in a 95°C water bath for 35 minutes. Samples were permeabilized in 0.2% Triton X-100 (Sigma-Aldrich) for 5min, blocked in 1.5% normal donkey serum (Jackson Immunoresearch) for 1 hour, and probed with primary antibodies against SOX2, PAX6, TBXT, NES, TUBB3, and CDH2 (Table S3) overnight at 4°C and secondary antibodies for 1 hour at room temperature. Nuclei were stained with a 1:10000 dilution of Hoechst 33342 (Thermo Fisher) included with secondary antibodies. Coverslips were mounted with anti-fade mounting medium (ProlongGold, Life Technologies) and samples were imaged on a Zeiss Axio Observer Z1 inverted microscope equipped with a Hamamatsu ORCA-Flash 4.0 camera.

### Whole Mount Lightsheet Imaging

4% paraformaldehyde-fixed paraffin-embedded samples (see “Histology, Immunocytochemistry, and Imaging”) were permeabilized with 1.5% Triton X-100 (Sigma-Aldrich) for 1 hour, blocked in 5% normal donkey serum (Jackson Immunoresearch) for 1 hour, and probed with primary and secondary antibodies (TableS3) overnight. Nuclei were stained with a 1:10000 dilution of Hoechst 33342 (Thermo Fisher) included with secondary antibodies. Samples were then embedded in 1.5% low melt agarose (BioReagent) and drawn up into ~1mm imaging capillaries and subsequently imaged on the Zeiss Z.1 Light sheet Microscope equipped with two PCO.edge SCMOS cameras at 5X and 20X (NA 1.34, aqueous objective).

### Flow cytometry

WTC and H1 cells were pretreated with either 2, 4, or 6 μM CHIR for 2 days, then dissociated from tissue culture plates with Accutase (STEMCELL Technologies) and washed with PBS. Similarly, the LBC-TBXT knockdown cells were pretreated with between 0 and 5 days of doxycycline concurrent with a final 2 days of 4 μM CHIR treatment, then dissociated and washed as described above. Cells were fixed for 20 minutes with 4% paraformaldehyde and washed 3x for 3 minutes with PBS. Samples were permeabilized in 0.5% Triton-X-100 (Sigma-Aldrich) for 30 minutes, then blocked in 1% normal donkey serum (Jackson Immunoresearch) for 1 hour, and probed with primary and secondary antibodies (TableS3) overnight. Samples were run on a LSR-II analyzer (BD Biosciences). Singlets were first identified by gating on forward scatter to side scatter ratio, then samples were gated into SOX2+/− and TBXT+/− using single stained controls for each cell line, then samples were assessed for percent SOX2+/TBXT+ cells. Analysis was conducted with a minimum of 20,000 events per sample.

### Bulk RNA Sequencing Sample and Library Preparation

Whole organoids differentiated in either 0μM CHIR or 4μM CHIR at days 1, 3, 5, 7, and 10 of the differentiation protocol (n=3 per condition per day) were lysed with RPE buffer with 5 mM 2-mercaptoethanol, and RNA was extracted using the RNeasy Mini Kit (Qiagen) and quantified using the NanoDrop 2000c (ThermoFisher Scientific). RNA-seq libraries were generated using the SMARTer Stranded Total RNA Sample Prep Kit (Takara Bio) and sequenced using NextSeq500/550 High Output v2.5 kit to a minimum depth of 25 million reads per sample. The sequences were aligned to GRCh37 using HiSat2^49^, reads were quantified using the featureCounts tool in the subread package^50^, and differential expression between 0μM and 4μM CHIR-treated organoids was assessed at each day using the limma/voom differential expression pipeline^51^. Longitudinal differential expression was assessed for each CHIR condition using the “l” normalization algorithm^52^. Raw data is available at Geo under the accession number GSE155382.

### Single Cell RNA Sequencing Sample and Library Preparation

Multiple organoid samples were combined and processed together using the MULTI-Seq technology^53^. Organoids were singularized using Accutase (STEMCELL Technologies) and washed with cold PBS. Cells were resuspended in PBS with lipid-modified Anchor and Barcode oligonucleotides (gift from Zev Gartner, UCSF) and incubated on ice for 5 minutes. A co-Anchor oligo was then added in order to stabilize membrane retention of the barcodes incubated for an additional 5 minutes on ice. Excess lipid-modified oligos were quenched with 1% BSA in PBS, washed with cold 1% BSA solution, and counted using a Countess II FL (Life Technologies). Single cell GEMs and subsequent libraries were then prepared using the 10X Genomics Single Cell V2 protocol with an additional anchor specific primer during cDNA amplification to enrich barcode sequences. Short barcode sequences (approx. 65-100bp determined by Bioanalyzer) were purified from cDNA libraries with two sequential SPRI bead cleanups. Barcode library preparation was performed according to the KAPA HiFi Hotstart (Kapa Biosystems) protocol to functionalize with the P5 sequencing adapter and library-specific RPIX barcode. Purified ~173bp barcode fragments were isolated with another SPRI bead cleanup and validation by Bioanalyzer. Raw data is available at Geo under the accession number GSE155383.

The sample library was sequenced on an Illumina NovaSeq yielding an average of 41,112 reads per cell and 6,444 cells. The MULTI-Seq barcode library was sequenced on an Illumina NextSeq yielding an average of 9,882 reads per barcode and enabling sample assignment for 4,681 of 6,124 unique UMIs detected (76.4% recovery), using the demultiplexing code provided by the MULTI-Seq protocol^53^.

### Genome Annotation, RNA-seq Read Mapping, and Estimation of Gene and Isoform Expression

The sample library was aligned to the human GRCh38 reference genome using Cell Ranger v1.2.0 (10x Genomics). Gene expression levels were assessed using the Seurat v3.0.0 analysis pipeline^54^. First cells were removed with fewer than 200 detected genes, fewer than 1,000 total detected transcripts, or which had greater than 10% mitochondrial gene expression. Next, expression levels were log normalized, and the top 2,000 variable genes calculated using the VST algorithm. The top 20 principal components were used to group cells into 2 clusters using a resolution of 0.3. Finally, top markers were detected for each cluster by detecting the top differentially expressed genes between both clusters, where at least 25% of cells in the cluster expressed the gene and the gene was expressed at least 0.25 log2 fold-change different from the remaining population. Clusters and gene expression were visualized on a two-dimensional UMAP projection of the first 15 principal components.

### Cluster Analysis

To assign cluster identity, the top markers for each cluster were tested for GO term enrichment using the biological process “enrichGO” function in the R package “clusterProfiler” v3.12 ^55^. In addition, differentiation maturity in each cluster was assessed by examining expression level of panels of early neuroectoderm markers, proliferation markers, markers of neuron fate commitment, and markers of cell types present in neural tube formation and axial extension^56^. Finally, to assess anterior-posterior position of each, panels of HOX genes were examined to assign rough position of each cluster along the head-tail axis^10,57,58^.

### Quantification of EdU and PH3 Localization

Sections stained for DAPI, EdU and ph3 were segmented by detecting the peaks of DAPI staining for each section using non-maximum suppression. The exterior of each section was then segmented by grouping DAPI+ cells into spatially contiguous clusters consisting of at least 5,000 pixels and which touch the image border in less than 1% of their total area. The exterior contour was then calculated for each region and fit to an elliptical model by total least squares^59^. All detected cells were projected onto the ellipse major axis, which was then normalized by total length. Where more detections fell on the left half of the major axis, the coordinates were reversed such that the right half of each projected section always contained the majority of detections. Detected cells that corresponded to ph3 or EdU fluorescence at 20% or more above the background level outside of any section were counted as positive detections. A 15-bin histogram was then used to calculate percentage of projected cells for each day at each point along the semi-major axis, and then gaussian kernel density estimation was used to produce the empirical distribution. Linear and quadratic models were fit using ordinary least squares over the total population.

### RNAScope

In situ hybridization for HOXB1, HOXC6, HOXB9 (probe information in Table S4) was performed on sections of 4% paraformaldehyde-fixed paraffin-embedded samples (see “Histology, Immunocytochemistry, and Imaging”) using the RNAscope Multiplex Fluorescent Reagent Kit v2 (Advanced Cell Diagnostics) and following the protocol outlined in User Manual 323100-USM. Sections were imaged on a Zeiss Axio Observer Z1 inverted microscope equipped with a Hamamatsu ORCA-Flash 4.0 camera.

### Statistical Analysis

Each experiment was performed with at least three biological replicates. Multiple comparisons were used to compare multiple groups followed by unpaired T-tests (two tailed) between two groups subject to a post-hoc Bonferroni correction. In gene expression analysis, three replicates were used for each condition, and all gene expression was normalized to control wildtype populations followed by unpaired T-tests (two tailed). Significance was specified as P-values< 0.05 unless otherwise specified in figure legends. All error bars represent standard error of the mean (SEM) unless otherwise noted in the figure legend.

## Data Availability

The Bulk and single cell RNA-seq data generated from this study will be made available on the Gene Expression Omnibus upon acceptance. The data analysis code generated in the live imaging analysis studies will be made available via GitHub repository upon acceptance.

## Acknowledgements

We would like to thank the Gladstone Light Microscopy and Histology Core, the Gladstone Stem Cell Core, The Gladstone Flowcytometry Core and the Gladstone Graphics Team for their support and experimental expertise. Additionally, we would like to specifically thank Dr. Vaishaali Natarajan for her expertise and assistance with flow cytometry.

## Author Contributions

ARGL, DAJ, and TCM were responsible for the conceptualization of the project and designing experiments. Original organoid differentiation protocol was developed by JCB. Differentiations were conducted by ARGL, DAJ, NHE, EAB, and EAG. Single cell RNA sequencing preparation and analysis were performed by ARGL, DAJ, NHE, and MZK. Image analysis script was written and implemented by DAJ and ARGL. Histological sectioning and staining were conducted by DAJ, NHE, EAB, and EAG. Light sheet microscopy and analysis was conducted by ARGL, DAJ, and EAG. Quantitative PCR was done by ARGL, DAJ, NHE, EAB, and MZK. RNAscope sample preparation and analysis was performed by ARGL. Cell lines were generated by ARGL, DAJ, MZK, and NMC. ARGL and EAB prepared the figures with input from all co-authors. All co-authors contributed to writing the original manuscript.

## Supplementary Data

### Supplementary Figures and Figure Legends

**Figure S1:**
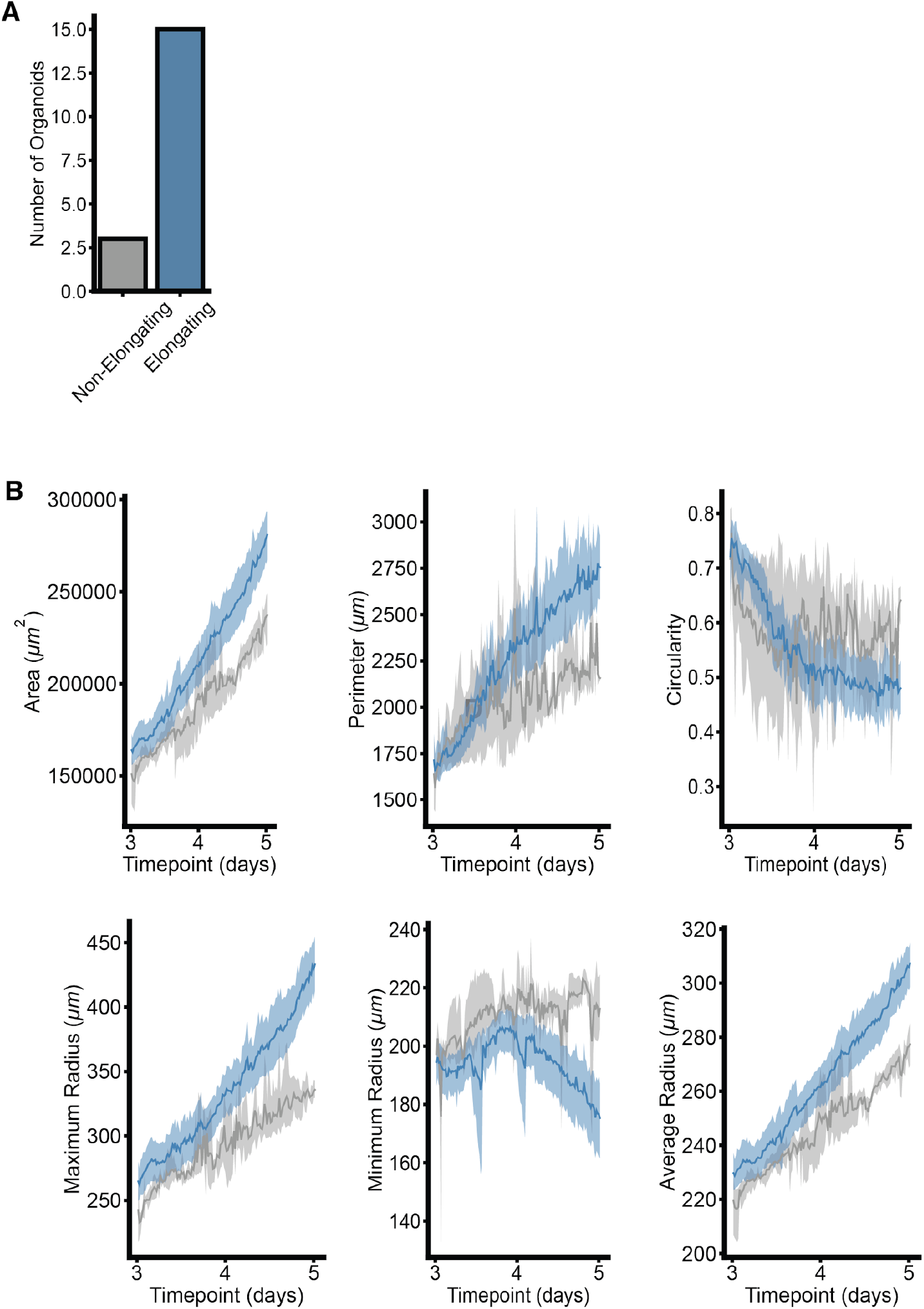
Time lapse imaging of organoid elongation. **A)** Number of extending verse non-extending aggregates in 4uM CHIR conditions (n=18). **B)** Quantification of area, perimeter, circularity, maximum, minimum, and average radius lengths of extending (blue) and non-extending (grey) aggregate videos. (mean value (dark line) with 95 percent confidence interval (light color)).

**Figure S2:**
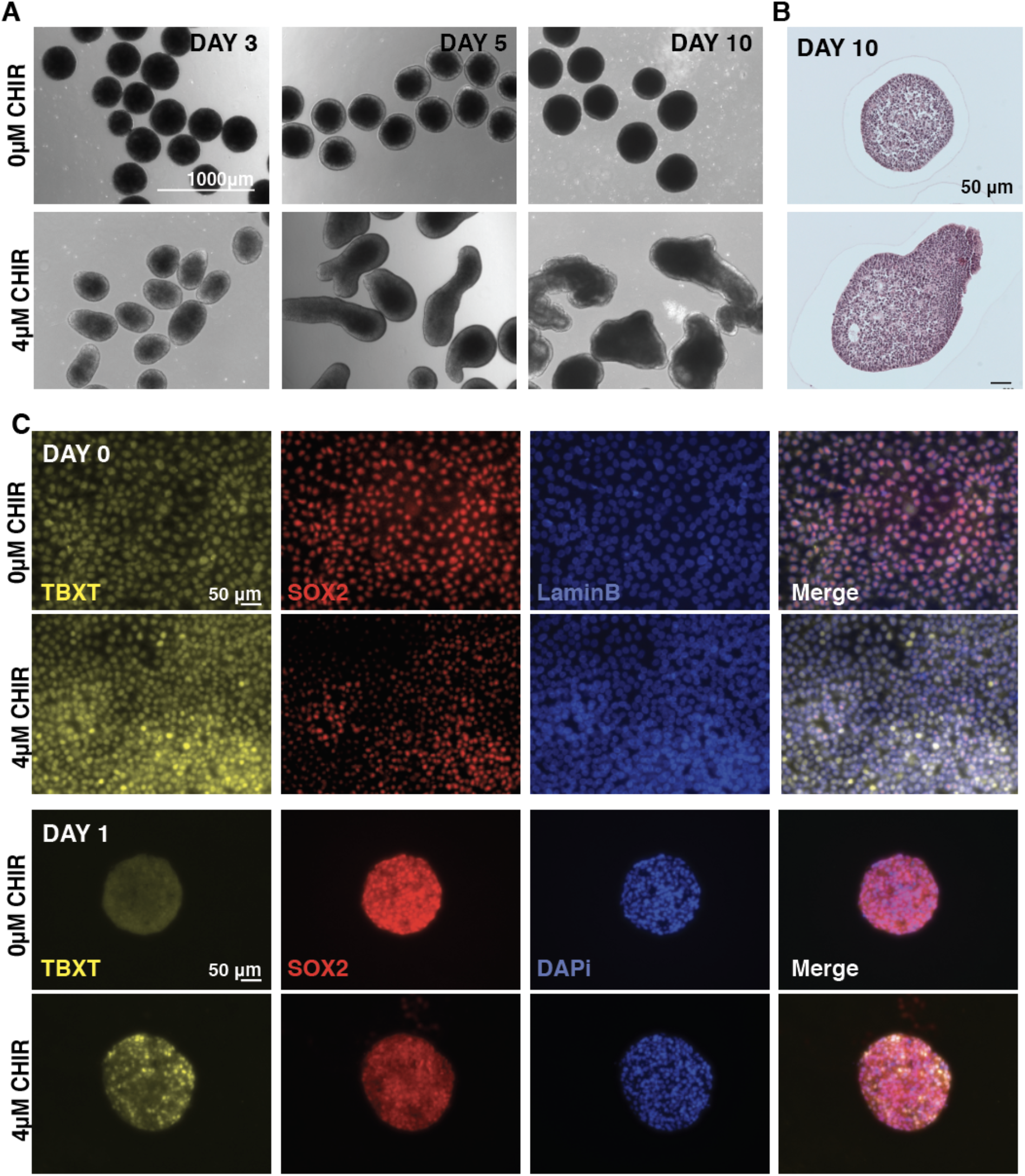
CHIR treatment results in extensions and increase of TBXT expression. **A)** Stereoscope images of differentiation performed with and without CHIR in the WTC hiPSC line. **B)** Histological sections of WTC organoids exposed to 0uM CHIR and 4uM CHIR **C)** Immunofluorescence images of TBXT and SOX2 in CHIR-treated and non-treated organoids.

**Figure S3:**
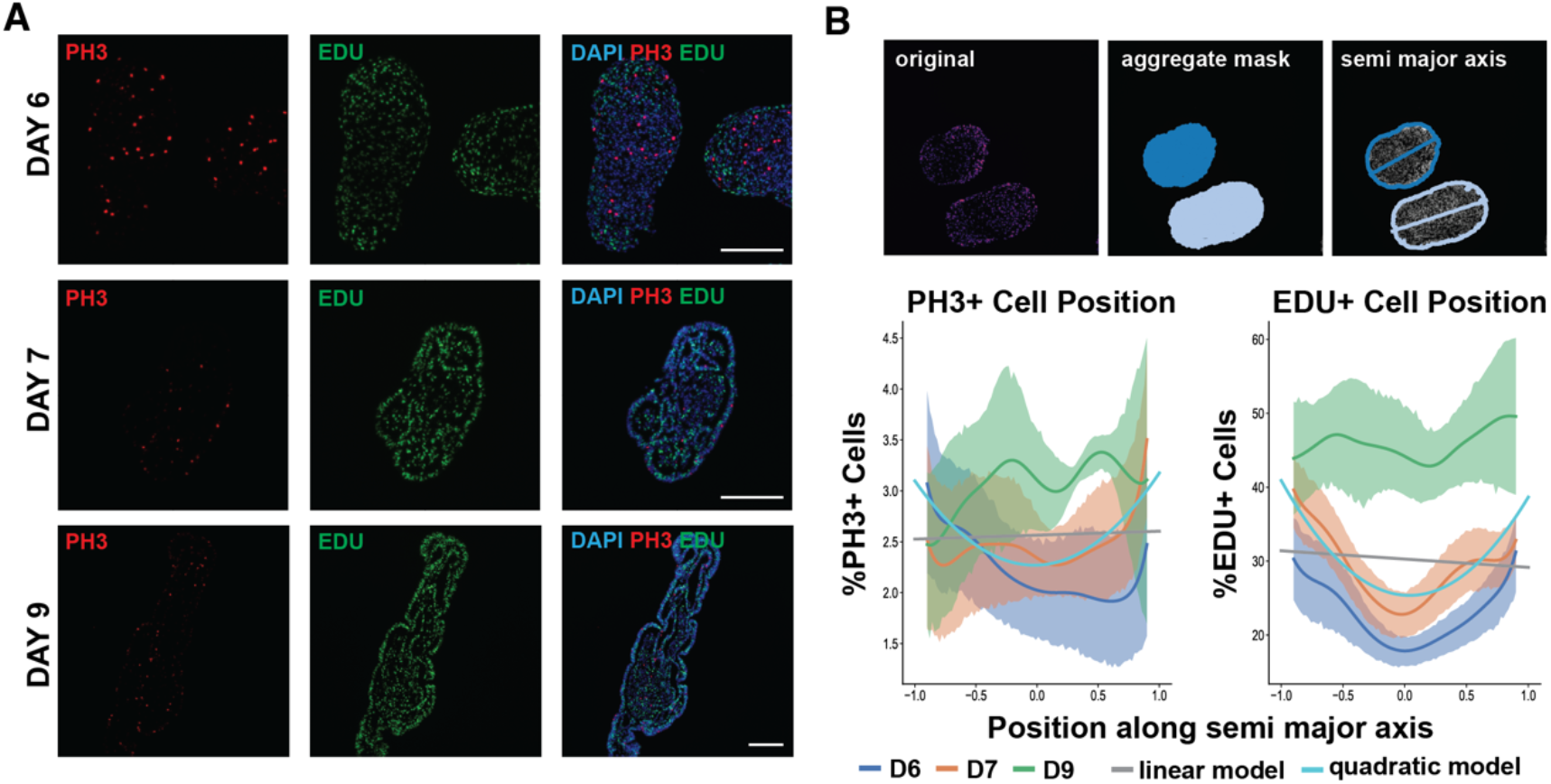
Cell proliferation occurs throughout the extending organoid. **A)** Immunofluorescence images of EdU incorporation and phospho-histone 3 (PH3) localization in extending organoids taken from paraffin sections. **B)** TOP: Segmentation of organoid sections showing LEFT) ph3 staining, MIDDLE) segmentation of nuclei by DAPI staining, RIGHT) a total least squares fit of the segmentation contour with the major axis superimposed. BOTTOM: Lineplots for PH3 or EdU expression along the normalized semi-major axis of aggregates at day 6, 7, and 9. Linear and quadratic least squares fits of PH3 and EdU expression along the normalized semi-major axis for all days.

**Figure S4:**
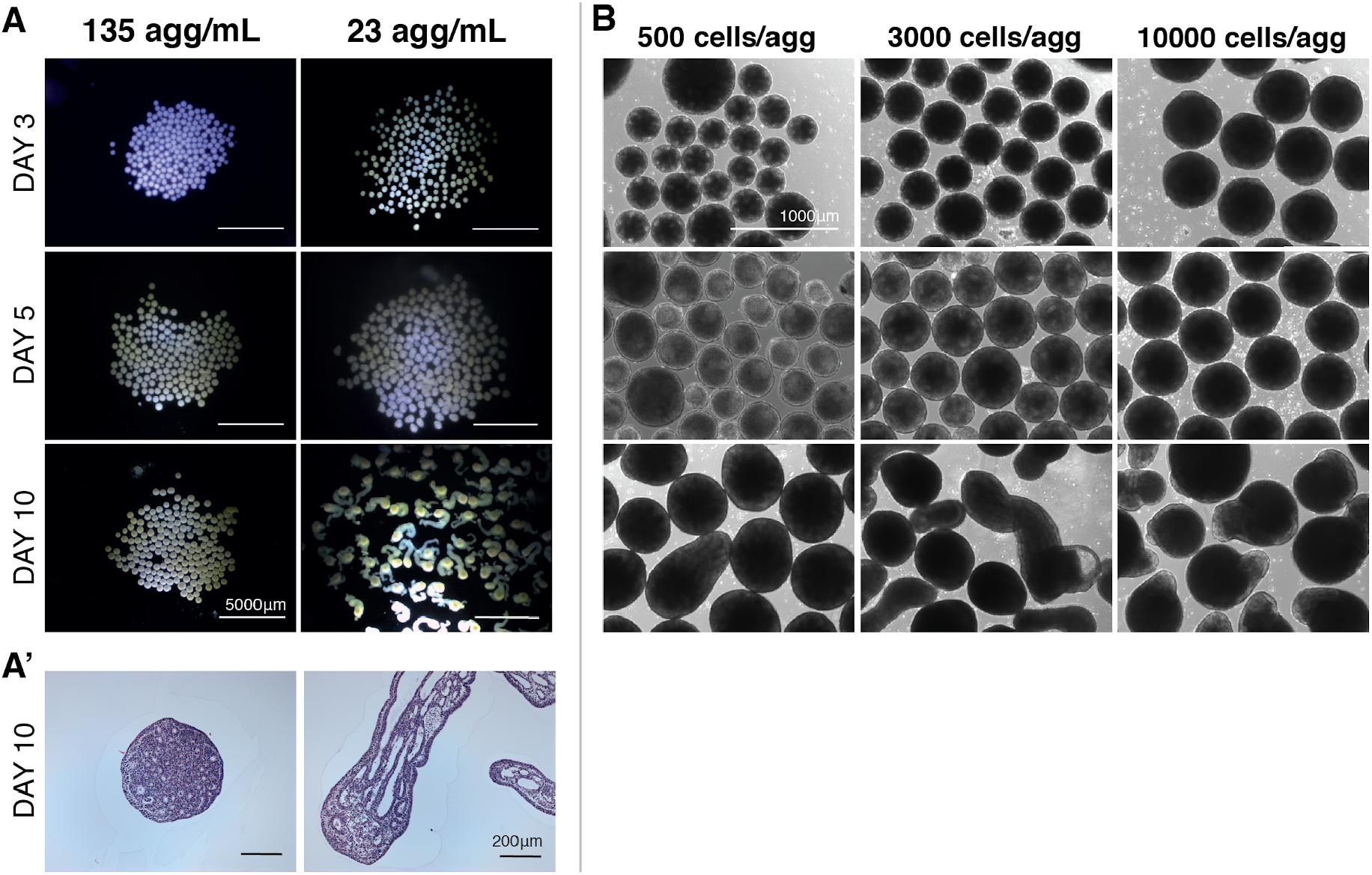
Organoid culture density changes morphology of extensions. **A)** Stereoscope images of organoids grown at high (135 aggregates/ml) or low (23 aggregates per ml) densities. **A’)** Histological stains on paraffin sections of organoids at day 10 of differentiation in high (left) vs low (right) density culture. **B)** Differentiations conducted in the WTC hiPSC line starting with different aggregate sizes.

**Figure S5:**
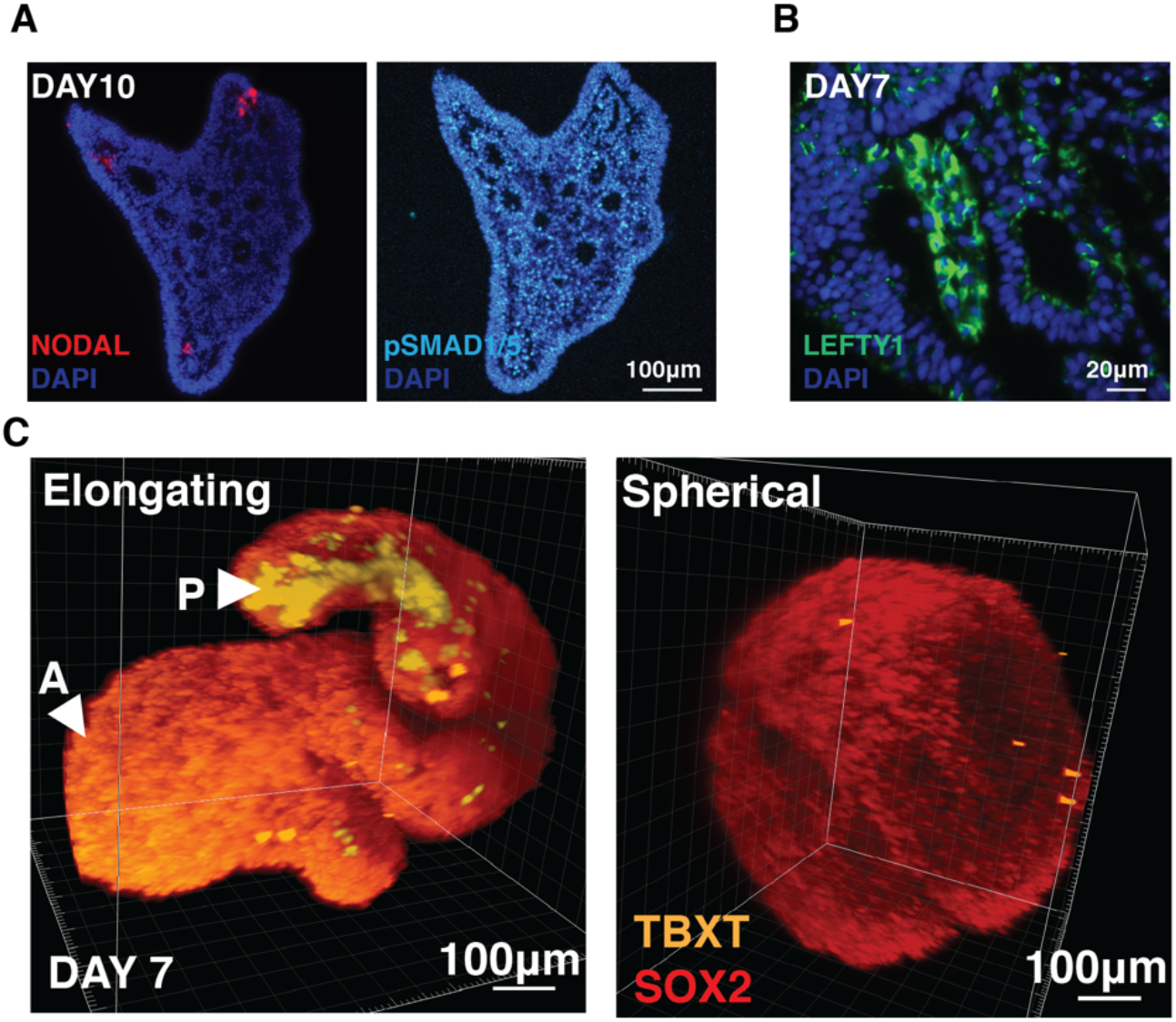
Gene expression distribution within extensions. **A)** Immunofluorescence staining of an extending organoid at day 10 of differentiation for NODAL and pSMAD1/5. **B)** Immunofluorescence staining of an extending organoid at day 7 of differentiation for LEFTY1. **C)** 3D reconstructions of light-sheet microscopy images of day 7 extending and non-extending organoids stained for TBXT and SOX2

**Figure S6:**
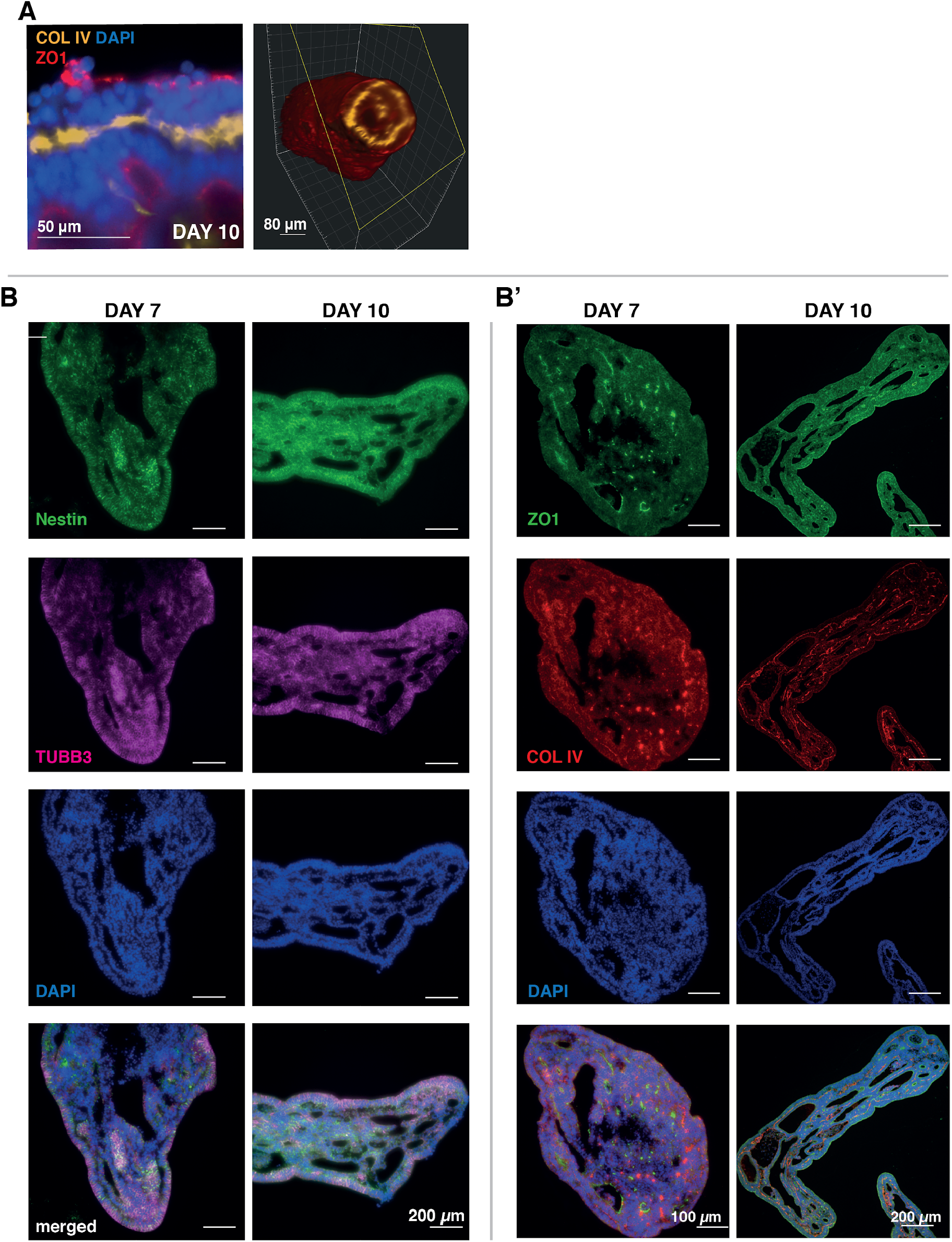
Apical-Basal polarity in extending organoids over time. B optical section from light-sheet imaging of extending stained for COL IV and ZO1 (left). 3D reconstruction showing internal ring of basement membrane within the posterior extensions (right). **B)** Immunofluorescence of paraffin sections of extending organoids at day 7 and day 10 of differentiation. **B’)** Immunostaining of paraffin sections of extending organoids at day 7 and 10 of differentiation for markers of basement membrane (COL IV) and tight junctions (ZO1).

**Figure S7:**
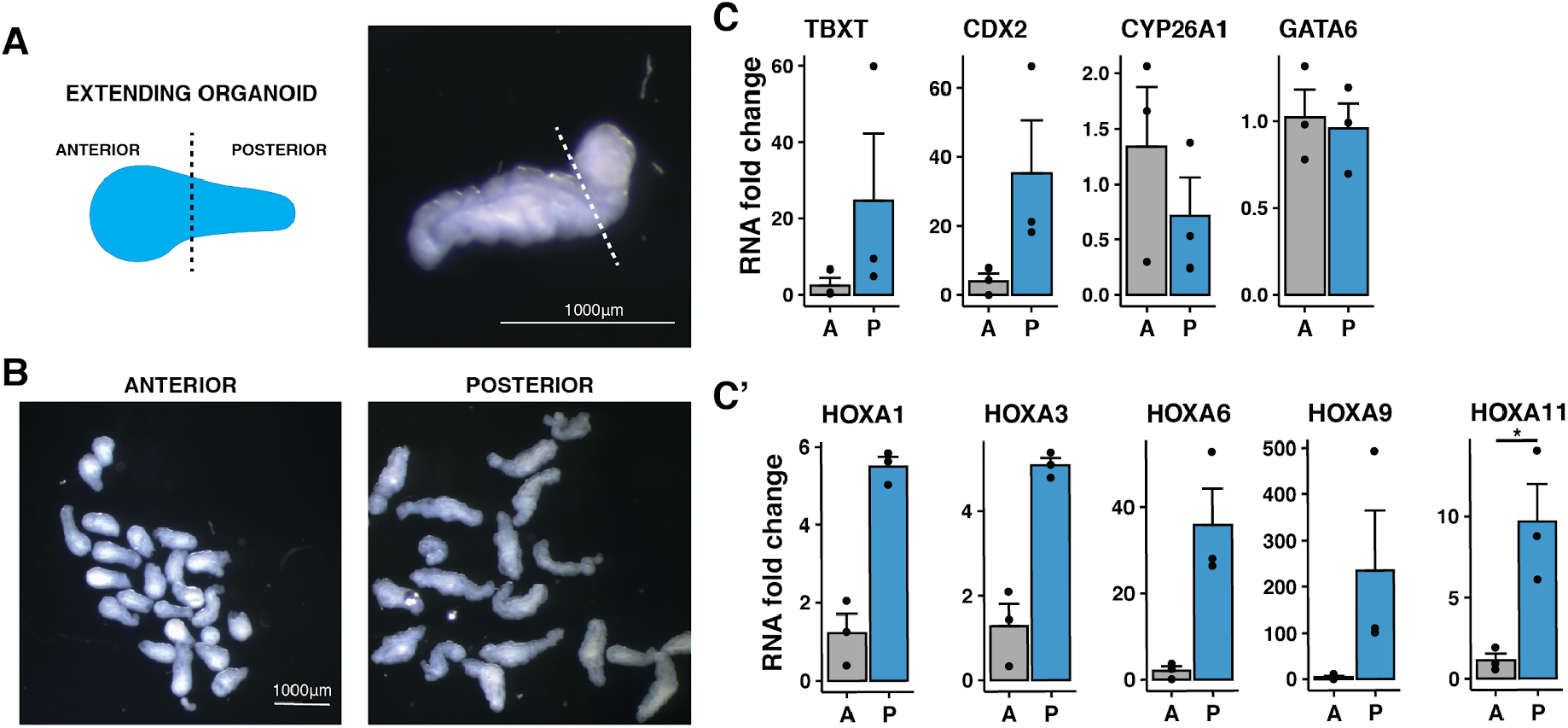
Dissections of extending organoids to isolate anterior and posterior regions. **A)** Schematic and example of anterior vs posterior regions. **B)** Stereoscope images of organoids after isolation of anterior or posterior region. **C-C’)** qPCR data of HOX genes and region specific genes in anterior vs. posterior of organoids (p < 0.05).

**Figure S8:**
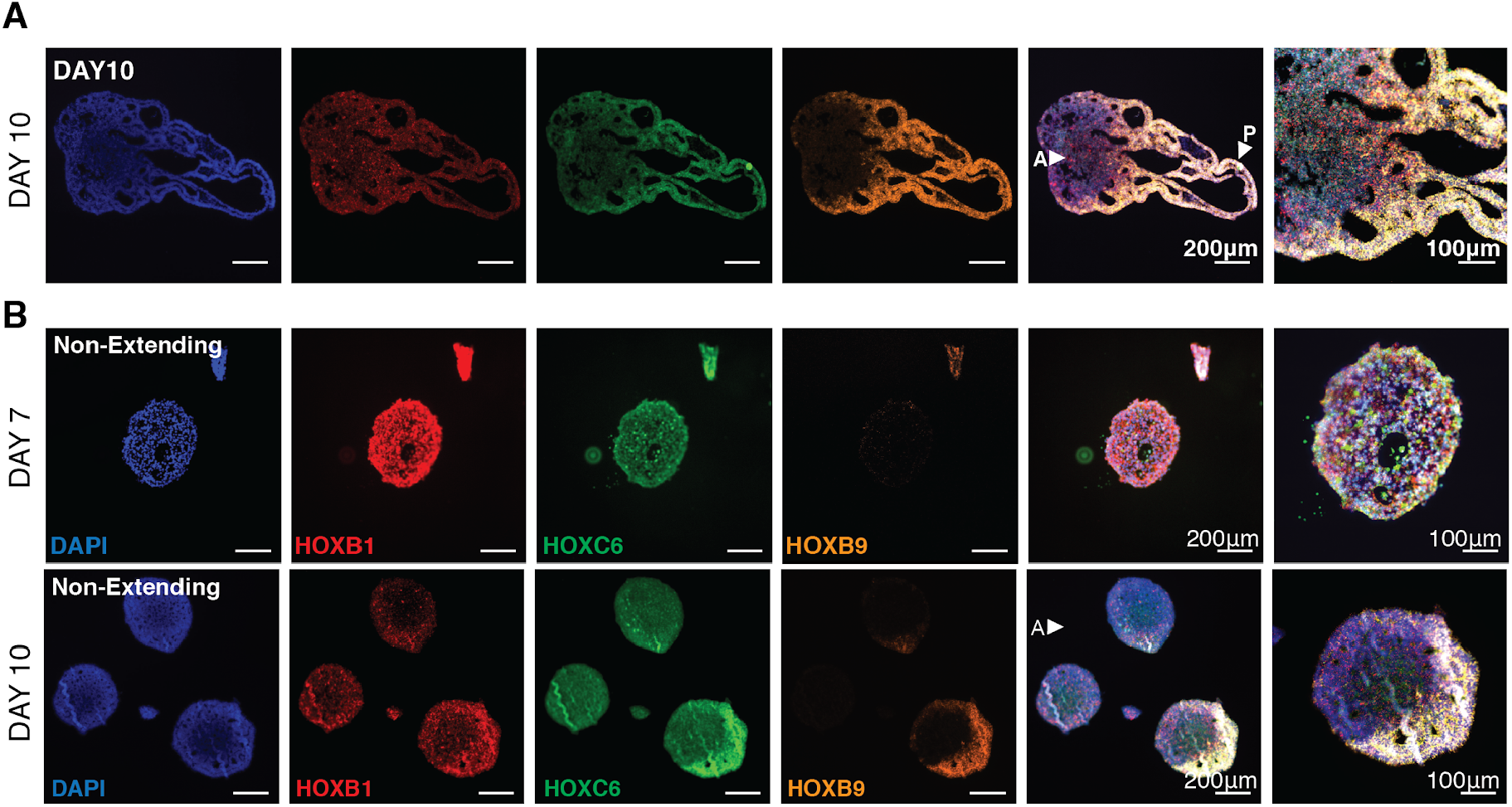
Limited segregation of HOX genes in non-extending organoids. **A)** RNAscope of sectioned extending organoids with probes for HOX genes marking different regions of the spine at day 10 of differentiation. **B)** RNAscope of sectioned non-extending organoids with probes for HOX genes marking different regions of the spine at day 7 and 10 of differentiation.

**Figure S9:**
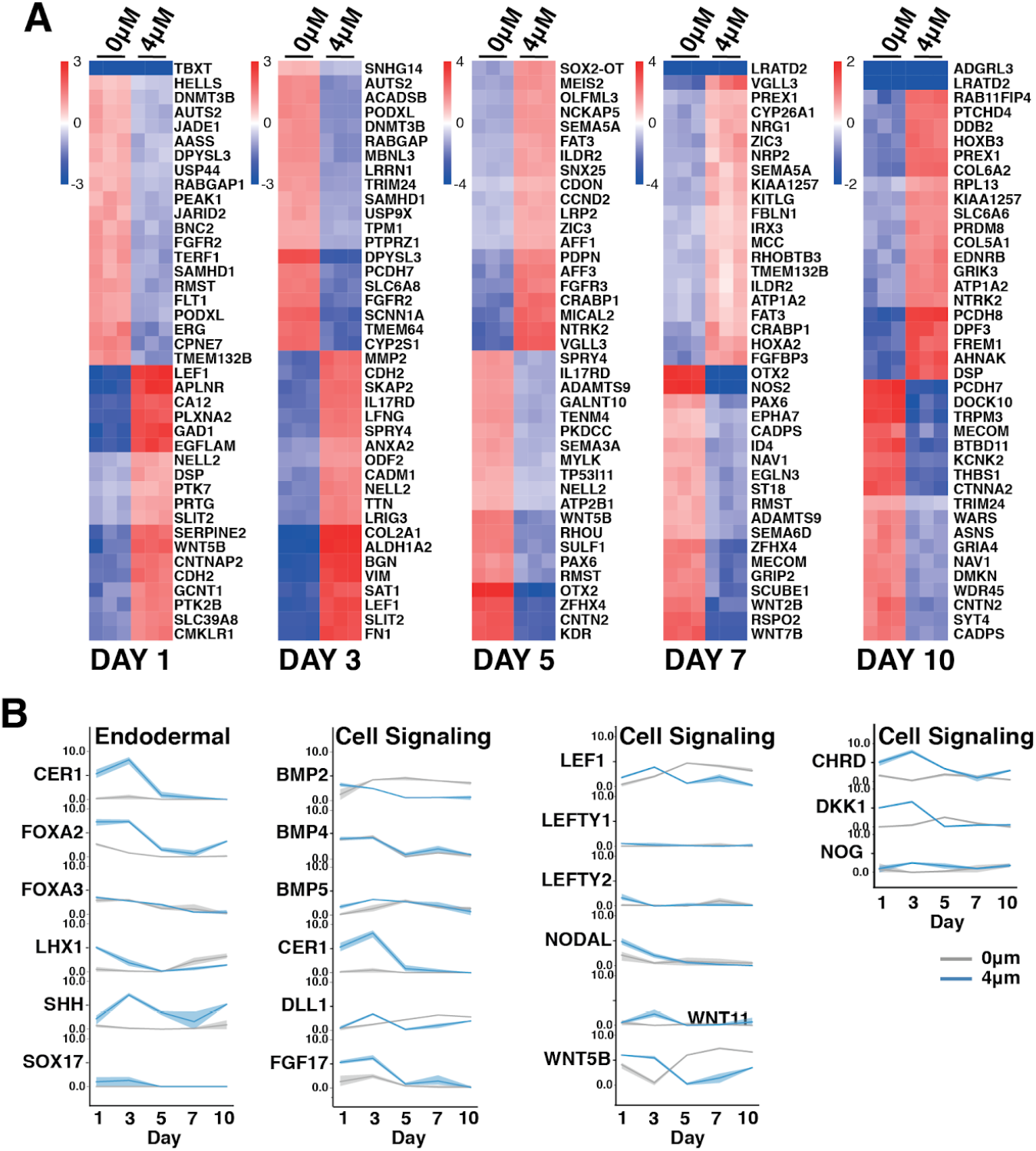
Bulk RNA sequencing of organoids with or without CHIR treatment. **A)** Top differentially expressed genes. **B)** Line plots showing gene expression of genes associated with signaling morphogens (grey line: 0μM CHIR; blue line: 4μM CHIR).

**Figure S10:**
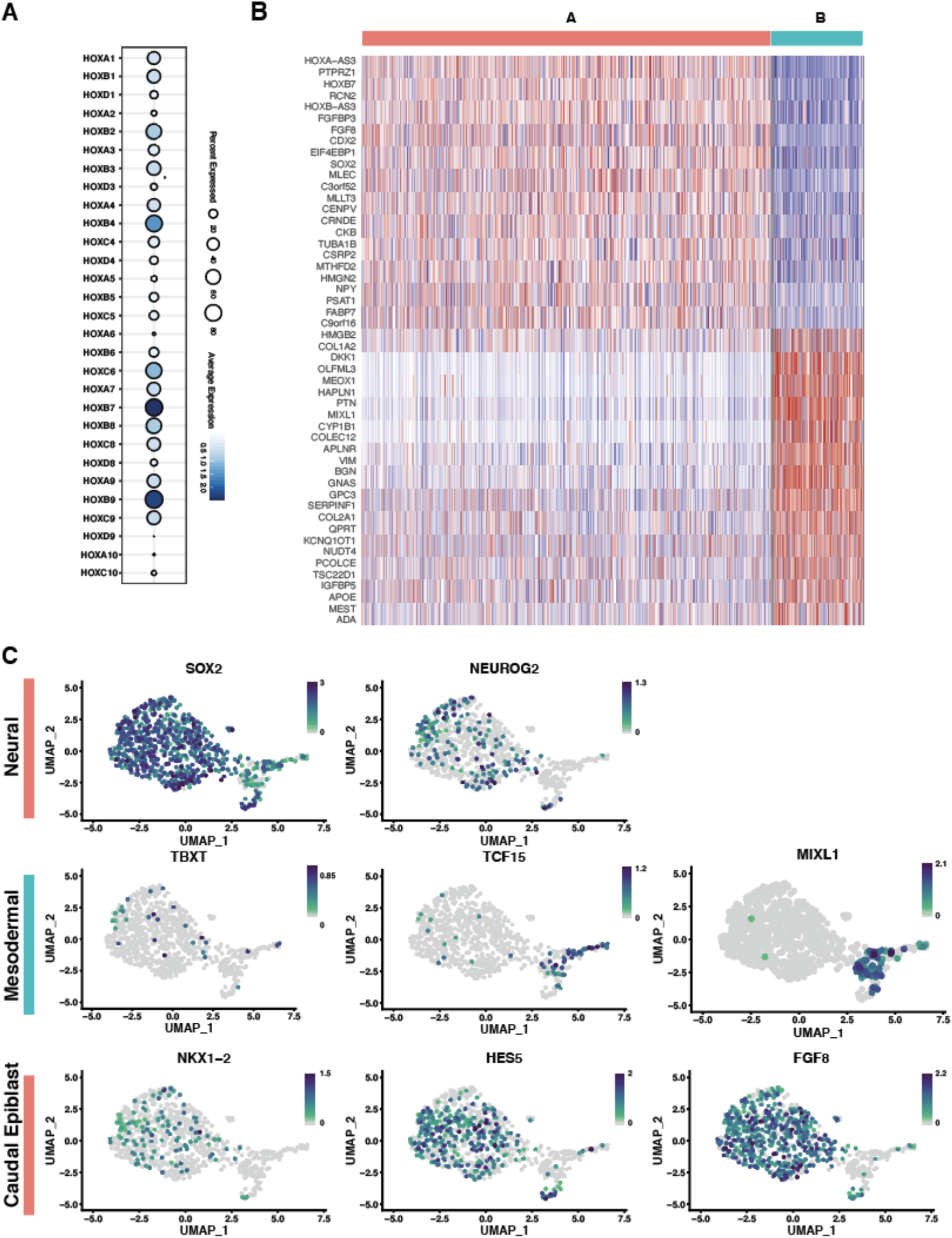
Single cell gene expression within extending organoids. **A)** Heat map of top differentially expressed genes between cluster A and B. **B)** Gene expression distribution plots. **C)** Dot plot of HOX gene expression within organoids.

**Figure S11:**
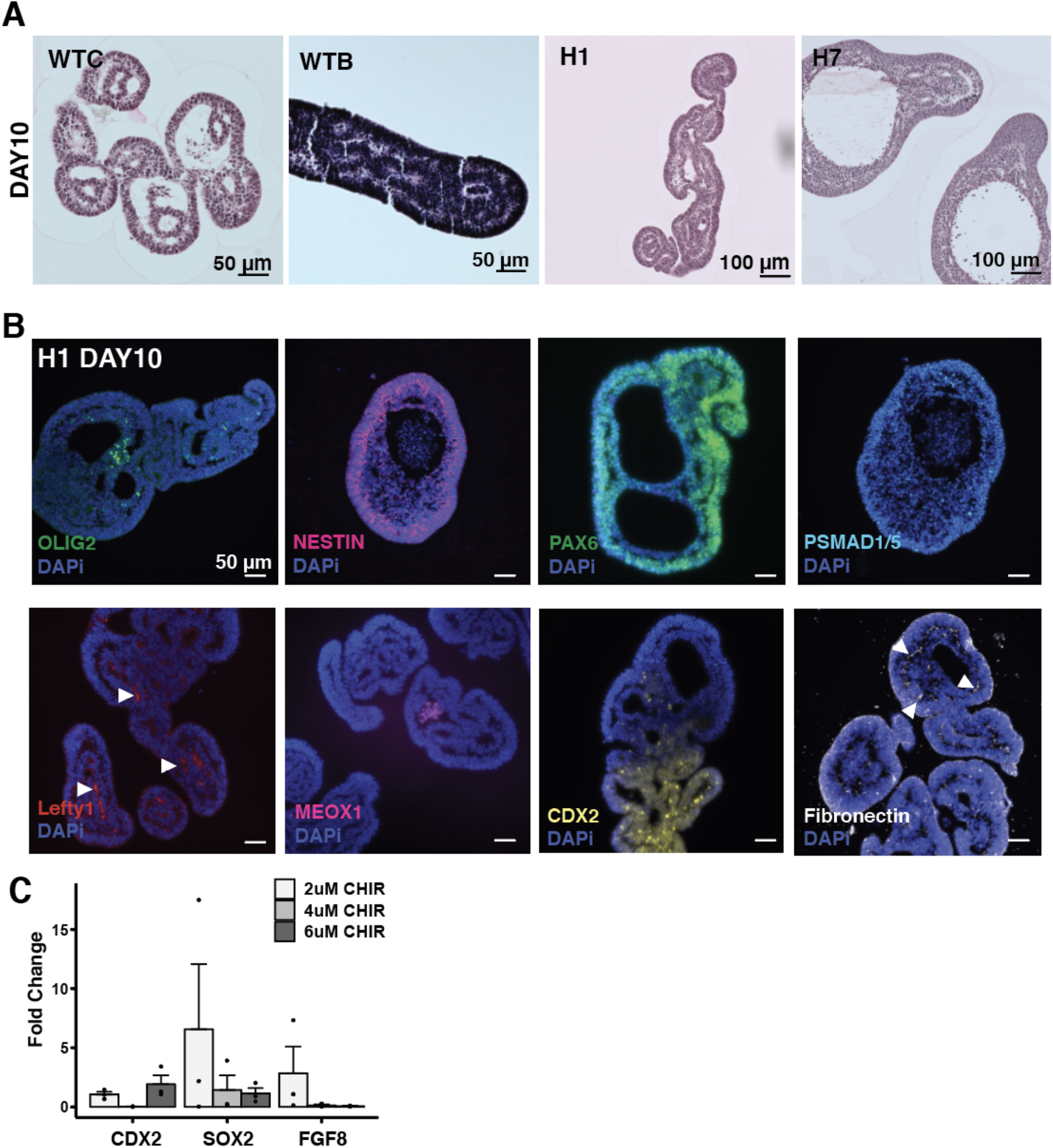
Extension reproducibility across hiPSC and hESC lines. **A)** Histology of the WTC and WTB hiPSC lines and H1 and H7 ESC line at extending (6uM) CHIR concentrations at day 10 of differentiation. **B)** Immunofluorescence staining of paraffin sections for OLIG2, Nestin, PAX6, pSMAD1/5, LEFTY1, MEOX1, CDX2, and Fibronectin at day 10 in H1 organoids (arrows indicate regions of positive staining). **C)** qPCR of caudal epiblast markers in the H1 ESC line across CHIR doses.

**Figure S12:**
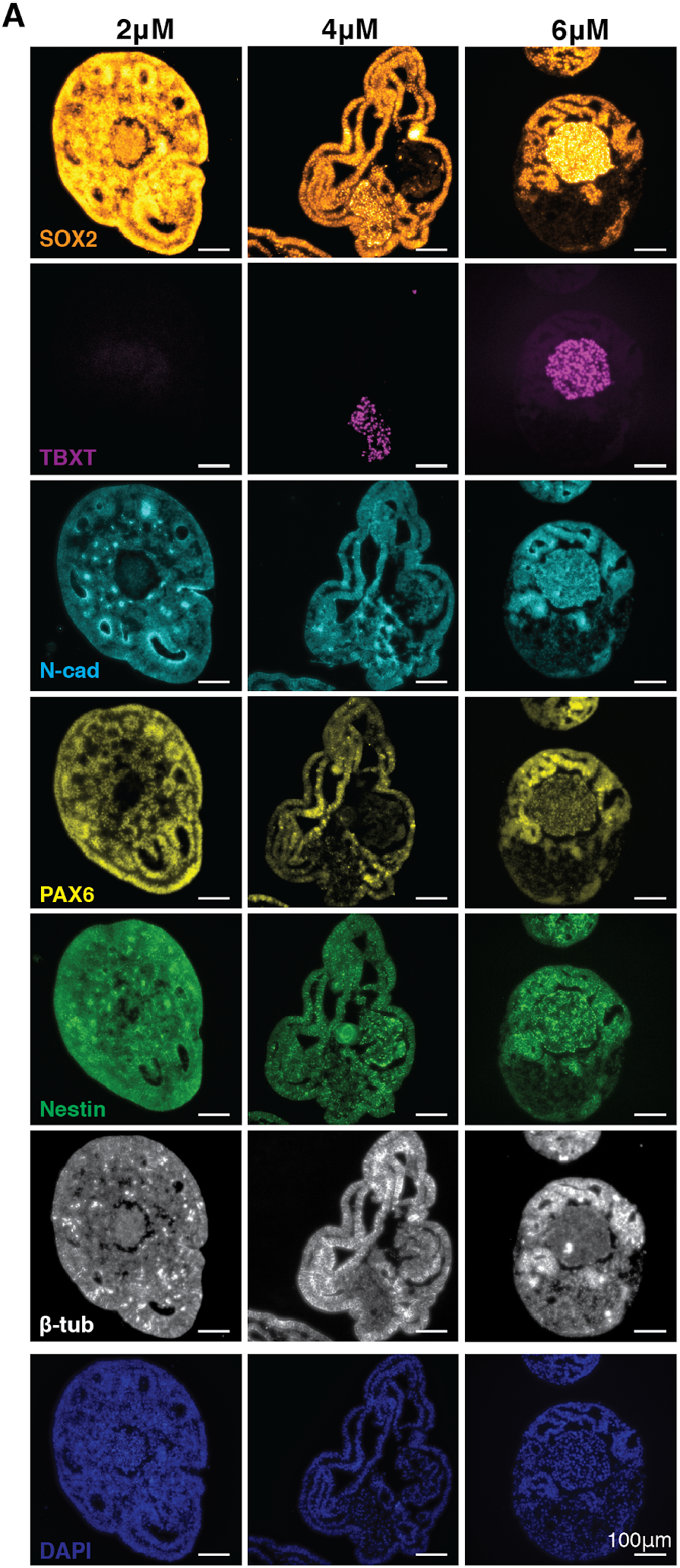
Neural cellular identities are maintained across CHIR dosage. **A)** Immunofluorescence of paraffin sectioned WTC hiPSC organoids at day 10 of differentiation from 2uM, 4uM, and 6uM CHIR concentrations.

**Figure S13:**
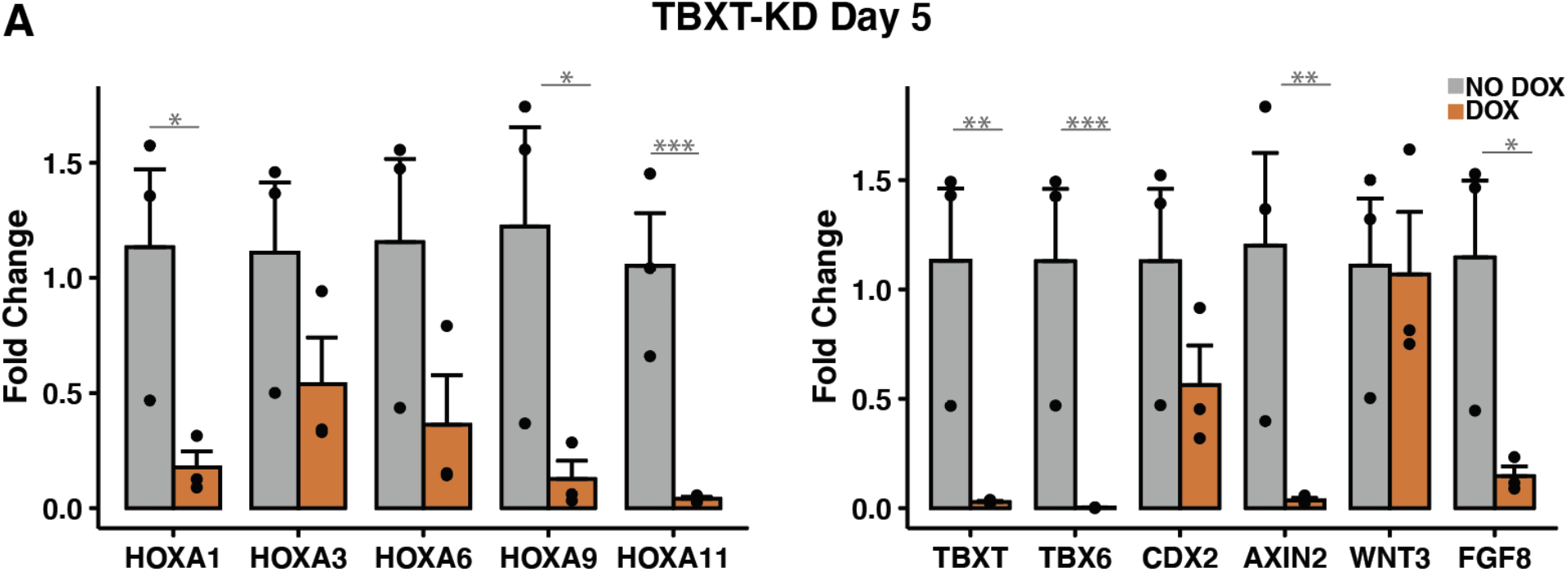
qPCR of TBXT KD at day 5 of differentiation. **A)** Quantification of gene expression by qPCR of HOX genes and genes associated with caudal fates in the TBXT knockdown organoids at day 5 of differentiation (* signifies p-value < 0.05, ** signifies p-value < 0.005, *** signifies p-value < 0.0005).

**Figure S14:**
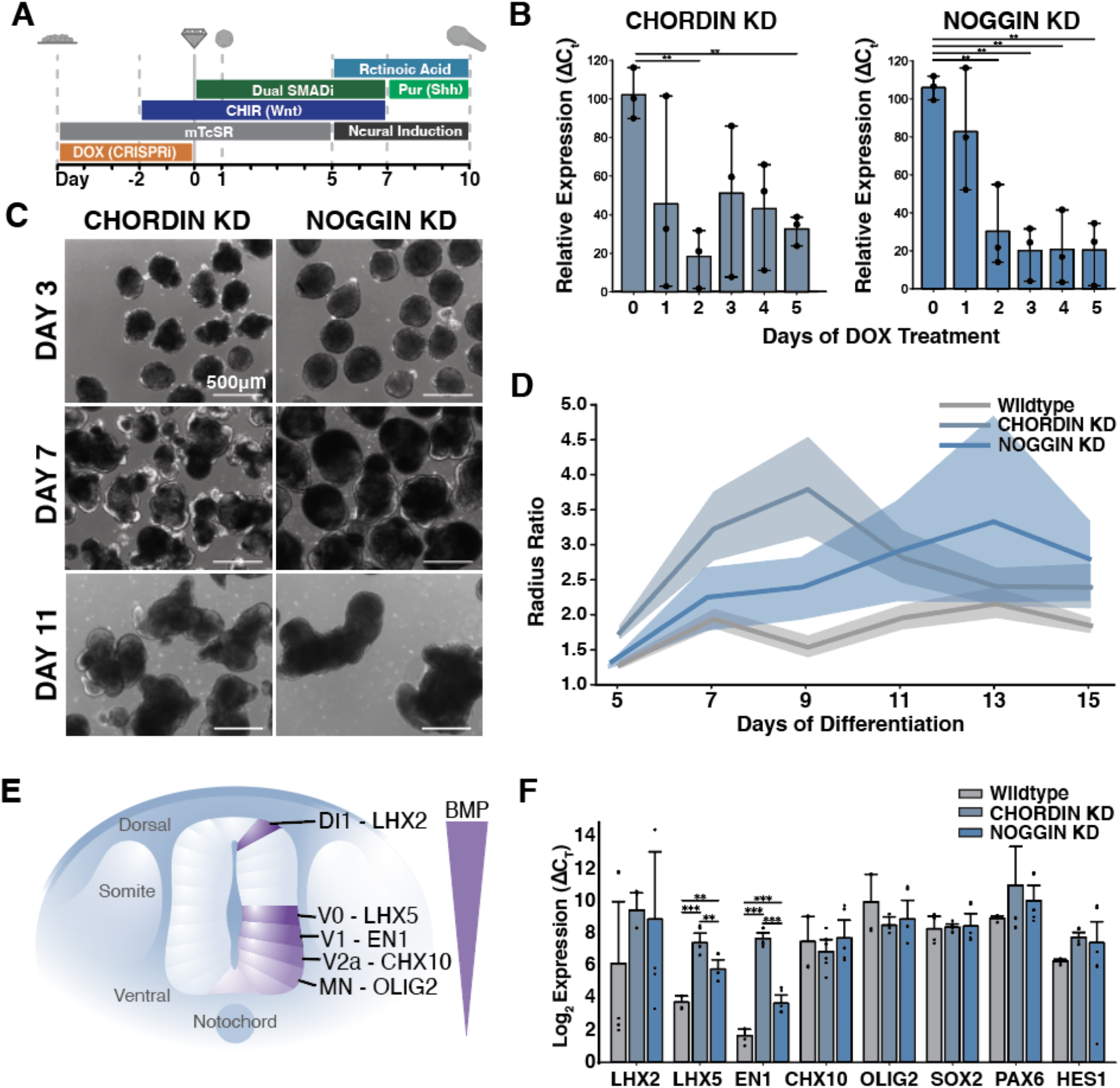
Manipulation of endogenous BMP signaling by CRISPR interference. **A)** Schematic of differentiation protocol timeline with knockdown induction. **B)** Quantification of Noggin and Chordin knockdown efficiency by reduction in mRNA expression levels measured by qPCR. **C)** Brightfield images of differentiation time course of knockdown hiPSC lines. **D)** Quantification of organoid length displayed as aspect ratio of major to minor axis in response to knockdown. **E)** Schematic depicting BMP signaling and progenitor specification within the developing neural tube. **F)** Quantification of mRNA expression of progenitor specific transcription factors by qPCR at day 17 of differentiation (n=3). Error bars depict standard deviations. Significance depicted by * indicating p<0.05, ** indicating p<.01, and *** indicating p<.001.

### Movie Legends

**Movie S1:** Elongating organoid dynamics. Movie of an elongating organoid cultured in a 96-well plate, taken over the course of differentiation from day 5 to day 7 with time steps every 20 minutes.

**Movie S2:** Non-elongating organoid dynamics. Movie of a non-elongating organoid cultured in a 96-well plate, taken over the course of differentiation from day 5 to day 7 with time steps every 20 minutes.

**Movie S3:** TBXT and SOX2 location in non-elongating organoid. Movie of a 3D rendered light-sheet image of a non-elongating organoid stained for TBXT (yellow) and SOX2 (red).

**Movie S4:** TBXT and SOX2 location in elongating organoid. Movie of a 3D rendered light-sheet image of an elongating organoid stained for TBXT (yellow) and SOX2 (red).

**Movie S5:** COL IV and ZO1 location in non-elongating organoid. Movie of a 3D rendered light-sheet image of a non-elongating organoid stained for COL IV (yellow), ZO1 (red), and DAPI (blue).

**Movie S6:** COL IV and ZO1 location in elongating organoid. Movie of a 3D rendered light-sheet image of an elongating organoid stained for COL IV (yellow), ZO1 (red), and DAPI (blue).

**Movie S7:** B III tubulin and Nestin location in non-elongating organoid. Movie of a 3D rendered light-sheet image of a non-elongating organoid stained for b III tubulin (TUBB; yellow) and Nestin (red).

**Movie S8:** B III tubulin and Nestin location in elongating organoid. Movie of a 3D rendered light-sheet image of an elongating organoid stained for b III tubulin (TUBB; yellow) and Nestin (red).

### Supplementary Tables

**Table S1:**
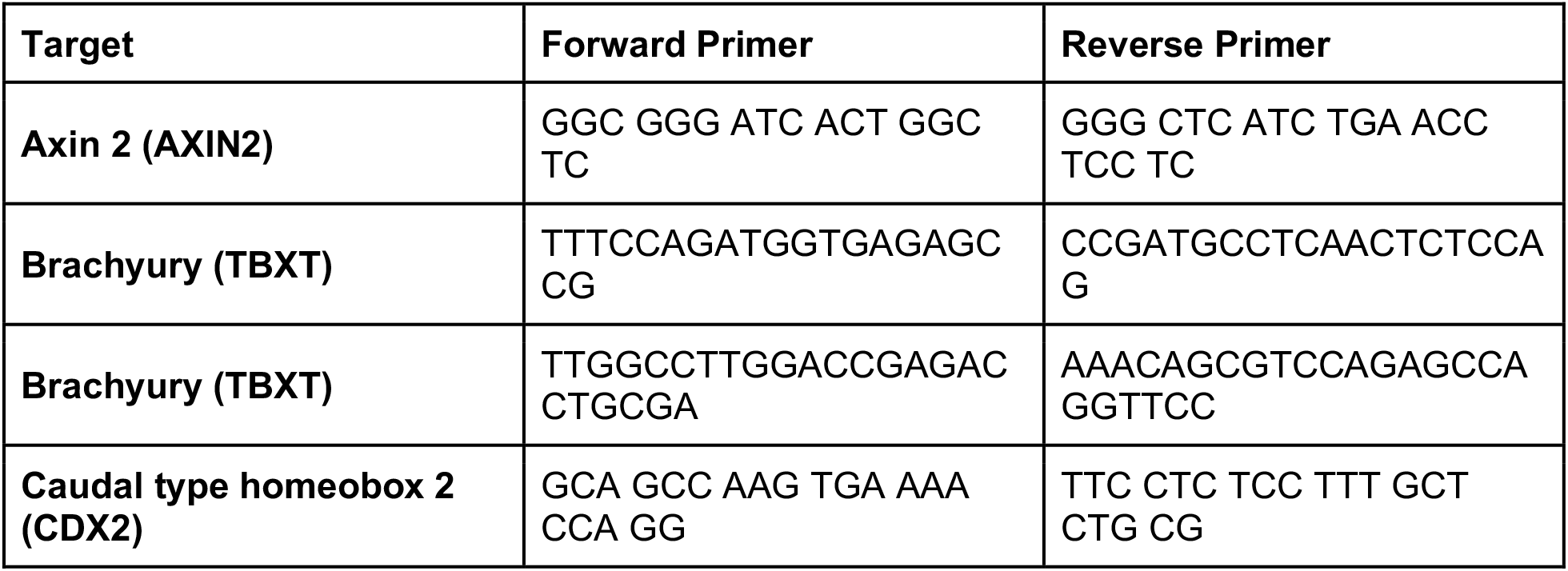
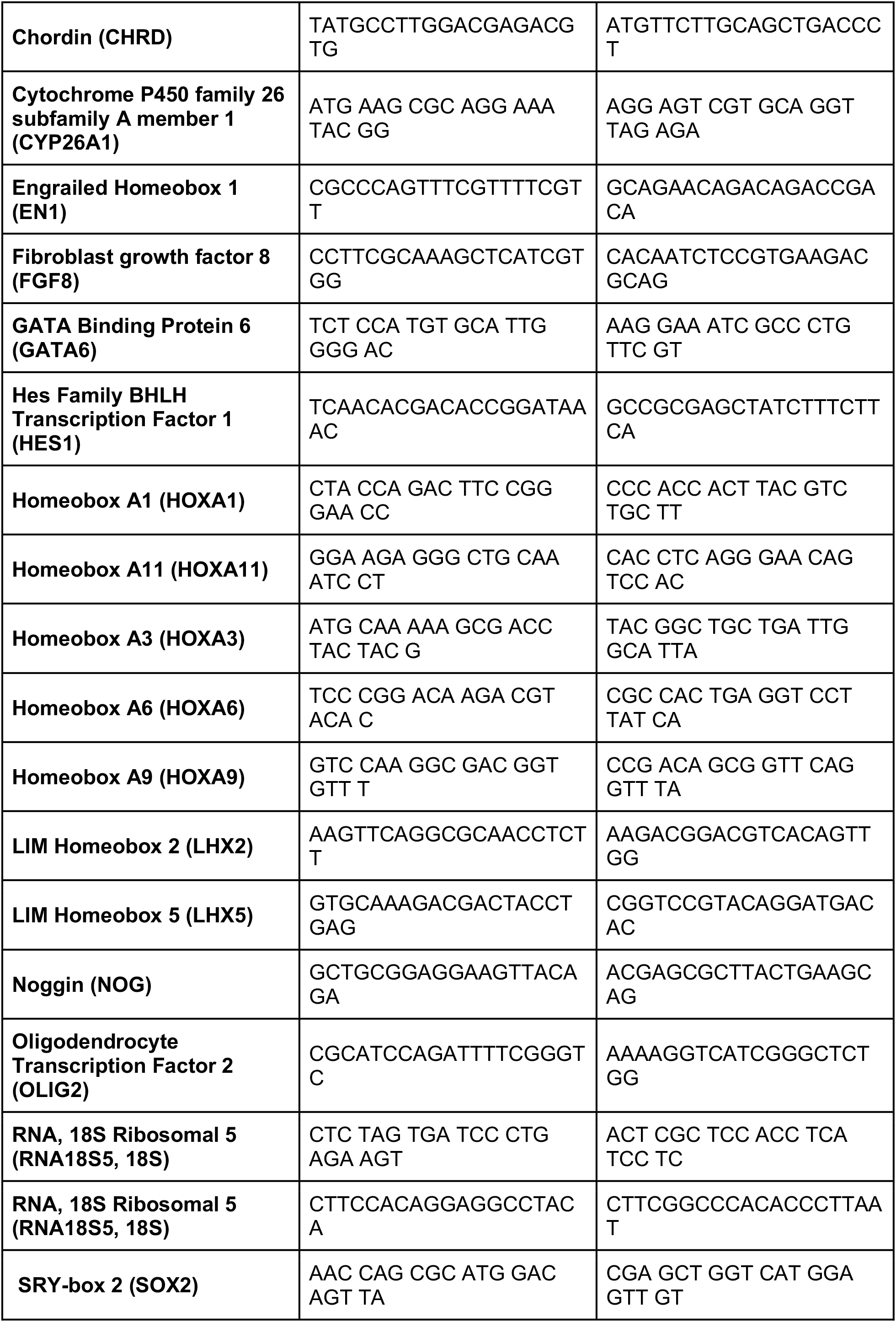
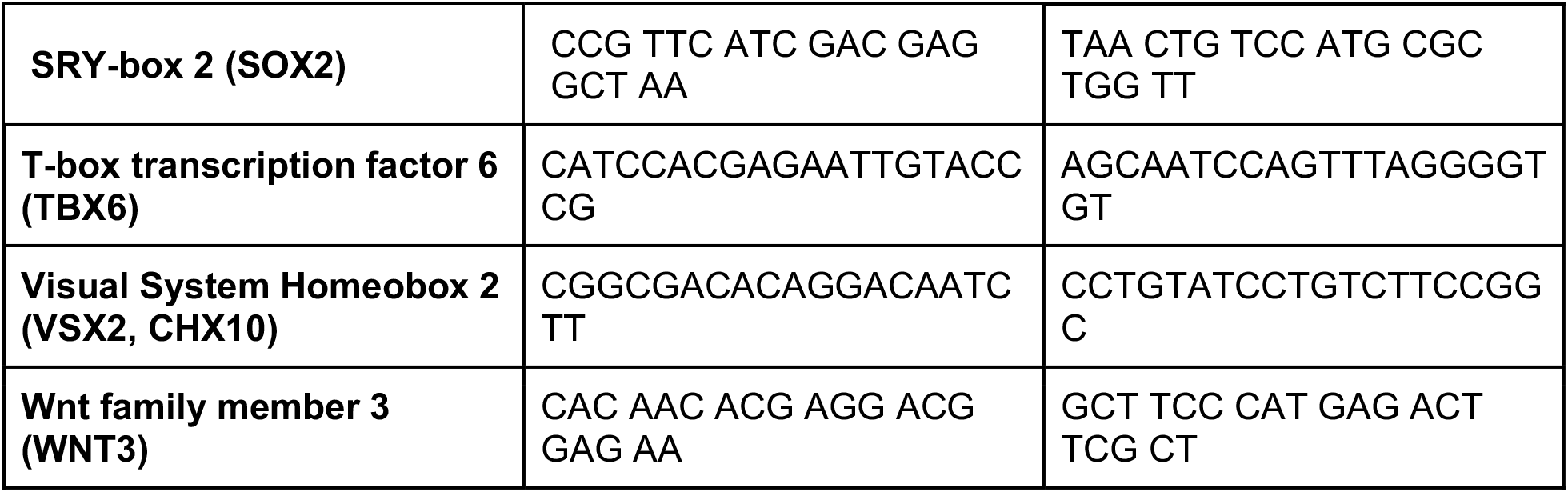
qPCR Primers.

**Table S2:**
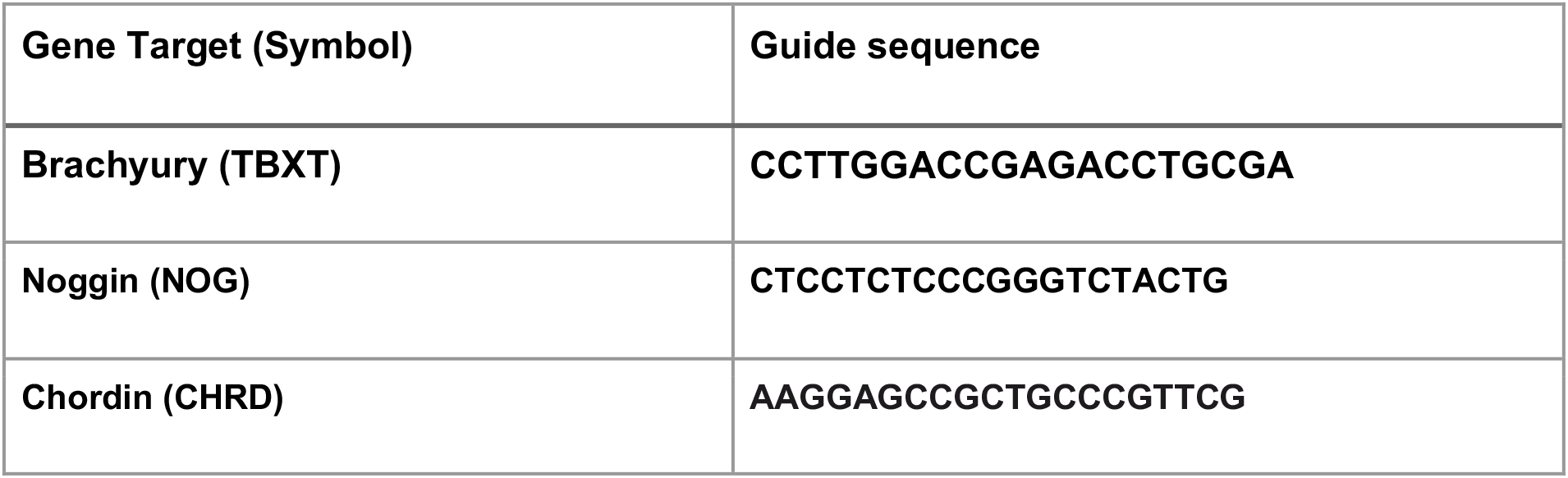
CRISPRi guides.

**Table S3:** Antibodies.

| **Gene Target (Symbol)** | **Species** | **Company (cat. #)** | **Dilution** |
| --- | --- | --- | --- |
| **SRY-box 2 (SOX2)** | mouse | Abcam (ab79351) | 1:400 |
| **Paired Box 6 (PAX6)** | rabbit | ThermoFisher Scientific (42-6600) | 1:400 |
| **Brachyury (TBXT)** | goat | ThermoFisher Scientific (PA5-46984) | 1:400, 1:300 |
| **Nestin (NES)** | mouse | Santa Cruz (SC-23927) | 1:400, 1:200 |
| **Hoescht DNA stain** | NA | ThermoFisher Scientific (62249) | 1:10000 |
| **Tubulin Beta 3 Class III (TUBB3/TUJ1)** | rabbit | Biolegend (802001) | 1:500 |
| **N-cadherin (CDH2)** | rabbit | Abcam (ab76057) | 1:400 |
| **Collagen type IV** | **goat** | **EMD Millipore (AB769)** | **1:75** |
| **Tight junction protein ZO-1 (TJP1/ZO1)** | **Mouse** | **Invitrogen (33-9100)** | **1:200** |
| **Caudal-Type Homeobox 2 (CDX2)** | **Goat** | **R&D Systems (AF-1979)** | **1:100** |
| **phospho-SMAD1/5 (pSMAD1/5)** | **Rabbit** | **Cell Signaling Technology (41D10)** | **1:800** |
| **Oligodendrocyte Transcription Factor-2 (OLIG2)** | **Rabbit** | **Abcam (ab9610)** | **1:400** |
| **Left-Right Determination Factor-1 (Lefty1)** | **Rabbit** | **ThermoFisher Scientific (PA5-19507)** | **1:1000** |
| **NODAL** | **Rabbit** | **ThermoFisher Scientific (PA5-23084)** | **1:200** |
| **Mesenchyme HomeoboxMEOX1** | **Mouse** | **ThermoFisher Scientific (TA-804716)** | **1:400** |
| **Fibronectin** | **Mouse** | **Sigma-Aldrich (F6140)** | **1:400** |
| **Phospho-Histone H3 (Ser10)** | **Rabbit** | **Cell Signaling Technology mAb #3377** | **1:1600** |

**Table S4:** RNAscope Probes.

| **Gene Target (Symbol)** | **channel** |
| --- | --- |
| **HOXB1** | **C2** |
| **HOXC6** | **C1** |
| **HOXB9** | **C3** |

## Main References

1 Steventon, B. et al. Species-specific contribution of volumetric growth and tissue convergence to posterior body elongation in vertebrates. Development 143, 1732–1741, doi:10.1242/dev.126375 (2016).

2 Wilson, V., Olivera-Martinez, I. & Storey, K. G. Stem cells, signals and vertebrate body axis extension. Development 136, 1591–1604, doi:10.1242/dev.021246 (2009).

3 Schiffmann, Y. Vol. 92 209–231 (Progress in Biophysics and Molecular Biology).

4 Cambray, N. & Wilson, V. Two distinct sources for a population of maturing axial progenitors. Development, 2829–2840, doi:10.1242/dev.02877 (2007).

5 Wymeersch, F. J. et al. Position-dependent plasticity of distinct progenitor types in the primitive streak. doi:doi:10.7554/eLife.10042 (2016).

6 Beddington, R. S. & Robertson, E. J. Axis development and early asymmetry in mammals. Cell 96, 195–209, doi:10.1016/s0092-8674(00)80560-7 (1999).

7 Liem, K. F., Tremml, G., Roelink, H. & Jessell, T. M. Dorsal Differentiation of Neural Plate Cells Induced by BMP-Mediated Signals from Epidermal Ectoderm. Cell 82, 969–979, doi:10.1016/0092-8674(95)90276-7 (1995).

8 Ericson, J. Graded Sonic Hedgehog Signaling and the Specification of Cell Fate in the Ventral Neural Tube. Cold Spring Harbor Symposia on Quantitative Biology 62, 451–466, doi:10.1101/SQB.1997.062.01.053 (1997).

9 McMahon, J. A. et al. Noggin-mediated antagonism of BMP signaling is required for growth and patterning of the neural tube and somite. Genes & Development 12, 1438–1452, doi:10.1101/gad.12.10.1438 (1998).

10 Diez del Corral, R. & Storey, K. G. Opposing FGF and retinoid pathways: a signalling switch that controls differentiation and patterning onset in the extending vertebrate body axis. Bioessays 26, 857–869, doi:10.1002/bies.20080 (2004).

11 Butts, J. C. et al. Differentiation of V2a interneurons from human pluripotent stem cells. Proc Natl Acad Sci U S A 114, 4969–4974, doi:10.1073/pnas.1608254114 (2017).

12 Gouti, M. et al. In vitro generation of neuromesodermal progenitors reveals distinct roles for wnt signalling in the specification of spinal cord and paraxial mesoderm identity. PLoS Biol 12, e1001937, doi:10.1371/journal.pbio.1001937 (2014).

13 Lancaster, M. A. & Knoblich, J. A. Organogenesis in a dish: Modeling development and disease using organoid technologies. doi:10.1126/science.1247125 (2014).

14 Turner, D. A., Rué, P., Mackenzie, J. P., Davies, E. & Arias, A. M. Brachyury cooperates with Wnt/β-catenin signalling to elicit primitive-streak-like behaviour in differentiating mouse embryonic stem cells. BMC Biology 12, 1–19, doi:doi:10.1186/s12915-014-0063-7 (2014).

15 Faustino Martins, J.-M. Self-Organizing 3D Human Trunk Neuromuscular Organoids. Cell Stem Cell 26, 172–186.e176, doi:10.1016/j.stem.2019.12.007 (2020).

16 Zheng, Y. et al. Dorsal-ventral patterned neural cyst from human pluripotent stem cells in a neurogenic niche. Science Advances 5, eaax5933, doi:10.1126/sciadv.aax5933 (2019).

17 Veenvliet, J. V. et al. Mouse embryonic stem cells self-organize into trunk-like structures with neural tube and somites. bioRxiv, doi:10.1101/2020.03.04.974949 (2020).

18 Beccari, L. et al. Multi-axial self-organization properties of mouse embryonic stem cells into gastruloids. Nature 562, 272–276, doi:doi:10.1038/s41586-018-0578-0 (2018).

19 Brink, S. C. v. d. et al. Single-cell and spatial transcriptomics reveal somitogenesis in gastruloids. Nature, 1–5, doi:doi:10.1038/s41586-020-2024-3 (2020).

20 Moris, N. et al. An in vitro model of early anteroposterior organization during human development. Nature 582, 410–415, doi:doi:10.1038/s41586-020-2383-9 (2020).

21 Warmflash, A., Sorre, B., Etoc, F., Siggia, E. D. & Brivanlou, A. H. A method to recapitulate early embryonic spatial patterning in human embryonic stem cells. Nature Methods 11, 847–854, doi:doi:10.1038/nmeth.3016 (2014).

22 Ortmann, D. & Vallier, L. Variability of human pluripotent stem cell lines. Current Opinion in Genetics & Development 46, 179–185, doi:10.1016/j.gde.2017.07.004 (2017).

23 Yamaguchi, T. P. Heads or tails: Wnts and anterior–posterior patterning. Current Biology 11, doi:10.1016/S0960-9822(01)00417-1 (2001).

24 Henrique, D., Abranches, E., Verrier, L. & Storey, K. G. Neuromesodermal progenitors and the making of the spinal cord. Development 142, 2864–2875, doi:10.1242/dev.119768 (2015).

25 Hookway, T. A., Butts, J. C., Lee, E., Tang, H. & McDevitt, T. C. Aggregate formation and suspension culture of human pluripotent stem cells and differentiated progeny. Methods 101, 11–20, doi:10.1016/j.ymeth.2015.11.027 (2016).

26 Oginuma, M. et al. Intracellular pH controls WNT downstream of glycolysis in amniote embryos. Nature, 1–4, doi:doi:10.1038/s41586-020-2428-0 (2020).

27 Beck, F., Erler, T., Russell, A. & James, R. Expression of Cdx-2 in the mouse embryo and placenta: possible role in patterning of the extra-embryonic membranes. Developmental dynamics: an official publication of the American Association of Anatomists 204, doi:10.1002/aja.1002040302 (1995).

28 Amin, S. et al. Cdx and T Brachyury Co-activate Growth Signaling in the Embryonic Axial Progenitor Niche. Cell Reports 17, 3165–3177, doi:10.1016/j.celrep.2016.11.069 (2016).

29 Herrmann, B. G., Labeit, S., Poustka, A., King, T. R. & Lehrach, H. Cloning of the T gene required in mesoderm formation in the mouse. Nature 343, 617–622, doi:doi:10.1038/343617a0 (1990).

30 T, P. et al. In vivo knockdown of Brachyury results in skeletal defects and urorectal malformations resembling caudal regression syndrome. Developmental biology 372, doi:10.1016/j.ydbio.2012.09.003 (2012).

31 Wilson, V., Rashbass, P. & Beddington, R. S. Chimeric analysis of T (Brachyury) gene function. (1993).

32 Wilson, V. & Beddington, R. Expression of T Protein in the Primitive Streak Is Necessary and Sufficient for Posterior Mesoderm Movement and Somite Differentiation-82000566.pdf. Developmental Biology 192, 45–58 (1997).

33 Chesley, P. Development of the short-tailed mutant in the house mouse - Chesley - 1935 - Journal of Experimental Zoology - Wiley Online Library. Journal of Experimental Zoology, doi:10.1002/jez.1400700306 (1935).

34 Bachiller, D. et al. The role of chordin/Bmp signals in mammalian pharyngeal development and DiGeorge syndrome. doi:10.1242/dev.00581 (2003).

35 Libby, A. R. et al. Spatiotemporal mosaic self-patterning of pluripotent stem cells using CRISPR interference. Elife 7, doi:10.7554/eLife.36045 (2018).

36 Mandegar, M. A. et al. CRISPR Interference Efficiently Induces Specific and Reversible Gene Silencing in Human iPSCs. Cell Stem Cell 18, 541–553, doi:10.1016/j.stem.2016.01.022 (2016).

37 Larson, M. H. et al. CRISPR interference (CRISPRi) for sequence-specific control of gene expression. Nature Protocols 8, 2180–2196, doi:doi:10.1038/nprot.2013.132 (2013).

38 Lancaster, M. A. et al. Cerebral organoids model human brain development and microcephaly. Nature 501, 373–379, doi:doi:10.1038/nature12517 (2013).

39 Renner, M. et al. Self-organized developmental patterning and differentiation in cerebral organoids | The EMBO Journal. The EMBO Journal, doi:10.15252/embj.201694700 (2020).

40 Sato, T. et al. Single Lgr5 stem cells build crypt-villus structures in vitro without a mesenchymal niche. Nature 459, 262–265, doi:doi:10.1038/nature07935 (2009).

41 Harrison, S. E., Sozen, B., Christodoulou, N., Kyprianou, C. & Zernicka-Goetz, M. Assembly of embryonic and extraembryonic stem cells to mimic embryogenesis in vitro. doi:10.1126/science.aal1810 (2017).

42 Trivedi, V. et al. Self-organised symmetry breaking in zebrafish reveals feedback from morphogenesis to pattern formation. bioRxiv, doi:10.1101/769257 (2019).

43 Marikawa, Y. et al., Exposure-based assessment of chemical teratogenicity using morphogenetic aggregates of human embryonic stem cells. Reproductive Toxicology 91, 74–91, doi:10.1016/j.reprotox.2019.10.004 (2020).

## Methods References

44 Miyaoka, Y. et al. Isolation of single-base genome-edited human iPS cells without antibiotic selection. Nature Methods 11, 291–293, doi:doi:10.1038/nmeth.2840 (2014).

45 Ludwig, T. E. et al. Feeder-independent culture of human embryonic stem cells. Nat Methods 3, 637–646, doi:10.1038/nmeth902 (2006).

46 Park, S., Kim, D., Jung, Y. G. & Roh, S. Thiazovivin, a Rho kinase inhibitor, improves stemness maintenance of embryo-derived stem-like cells under chemically defined culture conditions in cattle. Anim Reprod Sci 161, 47–57, doi:10.1016/j.anireprosci.2015.08.003 (2015).

47 Random Walks for Image Segmentation - IEEE Journals & Magazine, <https://ieeexplore.ieee.org/abstract/document/1704833> (of Publication: 25 September 2006).

48 Walt, S. v. d. et al. scikit-image: image processing in Python. PeerJ 2, doi:10.7717/peerj.453 (2014).

49 Kim, D., Langmead, B. & Salzberg, S. L. HISAT: a fast spliced aligner with low memory requirements. Nature Methods 12, 357–360, doi:doi:10.1038/nmeth.3317 (2015).

50 Liao, Y. et al. featureCounts: an efficient general purpose program for assigning sequence reads to genomic features. Bioinformatics 30, 923–930, doi:10.1093/bioinformatics/btt656 (2014).

51 Law, C. W., Chen, Y., Shi, W. & Smyth, G. K. voom: precision weights unlock linear model analysis tools for RNA-seq read counts. Genome Biology 15, 1–17, doi:doi:10.1186/gb-2014-15-2-r29 (2014).

52 Zhang, Z. et al. Novel Data Transformations for RNA-seq Differential Expression Analysis. Scientific Reports 9, 1–12, doi:doi:10.1038/s41598-019-41315-w (2019).

53 McGinnis, C. S. et al. MULTI-seq: sample multiplexing for single-cell RNA sequencing using lipid-tagged indices. Nature Methods 16, 619–626, doi:doi:10.1038/s41592-019-0433-8 (2019).

54 Butler, A., Hoffman, P., Smibert, P., Papalexi, E. & Satija, R. Integrating single-cell transcriptomic data across different conditions, technologies, and species. Nature Biotechnology 36, 411–420, doi:doi:10.1038/nbt.4096 (2018).

55 Yu, G., Wang, L. G., Han, Y. & He, Q. Y. clusterProfiler: an R package for comparing biological themes among gene clusters. Omics 16, 284–287, doi:10.1089/omi.2011.0118 (2012).

56 Tanabe, Y. & Jessell, T. M. Diversity and Pattern in the Developing Spinal Cord. doi:10.1126/science.274.5290.1115 (1996).

57 Bel-Vialar, S., Itasaki, N. & Krumlauf, R. Initiating Hox gene expression: in the early chick neural tube differential sensitivity to FGF and RA signaling subdivides the HoxB genes in two distinct groups. Development 129, 5103–5115 (2002).

58 Carpenter, E. M. Hox Genes and Spinal Cord Development. Developmental Neuroscience 24, 24–34, doi:10.1159/000064943 (2020).

59 Halir, R.; Flusser, J. “Numerically stable direct least squares fitting of ellipses”. In Proc. 6th International Conference in Central Europe on Computer Graphics and Visualization. WSCG (Vol. 98, pp. 125–132).

